# Host-encoded DNA methyltransferases modify the epigenome and host tropism of invading phages

**DOI:** 10.1101/2024.05.05.592563

**Authors:** Michiko Takahashi, Satoshi Hiraoka, Yuki Matsumoto, Rikako Shibagaki, Takako Ujihara, Hiromichi Maeda, Satoru Seo, Keizo Nagasaki, Hiroaki Takeuchi, Shigenobu Matsuzaki

## Abstract

Restriction modification (RM) systems are ubiquitous bacterial defense systems; however, some phages evade RM system and adapt to their bacterial hosts. In such cases, phages are thought to stochastically acquire DNA methylation from host-encoded DNA methyltransferases (MTases), facilitating host adaptation. However, no studies have directly compared the methylomes of host bacteria and their infecting phages. Here, we demonstrate the epigenetic landscape of adapted phages with diverse infection histories, focusing on the broad host-range phage KHP30T as its adapts to three *Helicobacter pylori* strains. Using a multistage infection system, we observed that the adapted phages displayed significantly high titers against the last infected *H. pylori* strain, suggesting an attendant change in host tropism. Single-molecule real-time sequencing revealed that methylated motifs were predominantly shared between the adapted phages and their most recent host. Our findings enhance our understanding of epigenetic phage–host interactions, which have significant implications for microbial ecology.

## INTRODUCTION

Epigenetic modifications regulate the activity of specific genes and cellular functions without altering DNA sequences. In eukaryotes, epigenetic modifications play a major role in various biological processes through modulating chromatin structure and organization.^1^ In particular, DNA methylation, catalyzed by methyltransferases (MTases), is a key epigenetic signal discovered early in studies of DNA as genetic material.^2,3^ In mammals, DNA methylation controls gene expression and chromatin structure, contributing to various cellular processes, including cell differentiation, genome imprinting, and X chromosome inactivation.^4^

As bacteria lack essential elements such as histones to structurally modify DNA, DNA methylation serves as a primary epigenetic regulator. ^2^ In prokaryotes, DNA methylation generally occurs in three forms: N6-methyladenine (m6A) and C5-methylcytosine (m5C), both also found in eukaryotes, and N4-methylcytosine (m4C), specific to bacteria and archaea.^2,5,6^ Prokaryotic DNA methylation regulates multiple biological processes, including cell cycle, gene expression, virulence, and host defense.^6^

Historically, one of the most well-studied prokaryotic systems involved in DNA methylation is the restriction modification (RM) system that defends against foreign DNA such as conjugative plasmids and phages.^7–9^ Most of these systems comprise an MTase and a restriction endonuclease (REase) that recognizes a specific target sequence for cleavage. These RM systems are classified into Types I, II, and III based on their subunit composition, cofactor requirements, recognition sequence structure, and mode of action.^10^ REases interact with their target DNA sequences to cleave the DNA, whereas MTases methylate these recognition sites to protect them from cleavage^.10^ Consequently, host genomic DNA is methylated by endogenous MTases and protected against REase digestion, whereas unmethylated foreign DNA is cleaved by REases. In contrast to these three major groups, Type IV systems consist solely of REase(s) that recognize and cleave methylated DNA.^10,11^

Methods for detecting DNA methylation have been particularly well developed for eukaryotic m5C, because the biological importance of m5C in mammalian cells over recent decades.^12^ However, these techniques are not directly applicable to bacterial methylomes, where m6A and m4C are more prevalent. Several approaches, such as restriction enzyme assays, and bisulfite sequencing, have been used to detect bacterial methylomes with higher prevalences of m6A and m4C; however, these methods have some limitations: restriction enzyme assays cannot detect unknown motifs, Sanger sequencing has low throughput, and bisulfite sequencing cannot detect m6A and m4C.^13^ Therefore, further development of DNA methylation detection methods applicable to bacterial methylomes is needed.

Recent advances in sequencing technologies, including single-molecule real-time (SMRT) sequencing have addressed this problem. SMRT sequencing was the first to enable simultaneous detection of all three major forms of prokaryotic DNA methylation, leading to extensive analysis of bacterial and archaeal methylomes and significant expansion of REBASE, a database of information about components of RM systems.^13–18^ In addition, methylome analysis has been extended to viruses and phages.^19–22^ Recent studies revealed that some DNA viruses encode MTases in their genomes and use them as part of their replication strategy.^21,23,24^ In contrast, viruses that do not encode MTases can stochastically acquire DNA methylation from host-encoded MTases during their adaptation to the host strain. Experiments with microbes possessing well-characterized RM systems such as *Escherichia coli, Bacillus subtilis, Haemophilus* spp., and *Salmonella* spp. suggest that phage genomes are modified by host-encoded MTases during infection, allowing adapted phages to develop a “protective suit” against the host RM systems^.25^ However, no study has directly compared the methylomes of hosts and phages to determine whether adapted phages acquire DNA methylation from host-encoded

### MTases

Here, we demonstrate the epigenomic landscape of phages with distrinct histories of infection. To model bacteria-phage interactions mediated by host-encoded MTases, we established a multistage infection system and generated nine adapted phages with different infection histories using three strains of *Helicobacter pylori* and the *H. pylori* phage KHP30T. The adapted phages showed high infectious titers against the strains of *H. pylori* infected in the last generation but low titers against other strains. SMRT sequencing revealed that methylated motifs in the KHP30T genome were shared with those in the last-infected *H. pylori* strains. These findings suggest that the phage genome is modified by host-encoded MTases, resulting in the phage DNA evading cleavage by cognate REases. Our results provide direct evidence that adapted phages stochastically acquire DNA methylation from host bacteria and offer new insights relevant to phage therapy design.

## RESULTS

### Titration of adapted phages

We designed multistage infection systems using the KHP30T phage and three *H. pylori* strains (26695, 3401T, and HPK5) to obtain adapted phages with different infection histories (Figures 1 and S1). Using this strategy, nine adapted phages (K2, K23, K232, K23H, K23H3, K2H, K2H2, K2H3, and K2H3H) were retrieved. For example, K23 represents a <u>K</u>HP30T phage adapted to *H*. *pylori* <u>2</u>6695 in the first stage and then to *H. pylori* <u>3</u>401T in the second stage. Infectious titers were measured using a plaque assay. Adapted phages showed high titers against the *H. pylori* strain in the last generation of their adaptation but lower titers against non-adapted or previously adapted hosts (Figures 2 and S2). For example, K2, which was adapted to *H. pylori* 26695 in the first stage (Figure 1), showed a significantly higher titer (9.2 log10 plaque forming units [PFU]/mL) against 26695, but a markedly lower titer against 3401T and HPK5 (3.1 and 3.9 log10 PFU/mL, respectively) (*p* < 0.01, Tukey–Kramer test). Focusing on second-stage phages, K23 showed a significantly higher titer (8.9 log10 PFU/mL) against *H. pylori* 3401T than against 26695 and HPK5 (0.8 and 1.8 log10 PFU/mL, respectively) (*p* < 0.01), even though K23 had adapted to 26695 in the first stage. K2H also showed the same trend: a titer against *H. pylori* HPK5 (8.2 log10 PFU/mL) that was significantly higher than that against 26695 and 3401T (1.9 and 2.4 log10 PFU/mL, respectively, *p* < 0.01) (Figure 2).

**Figure 1.**
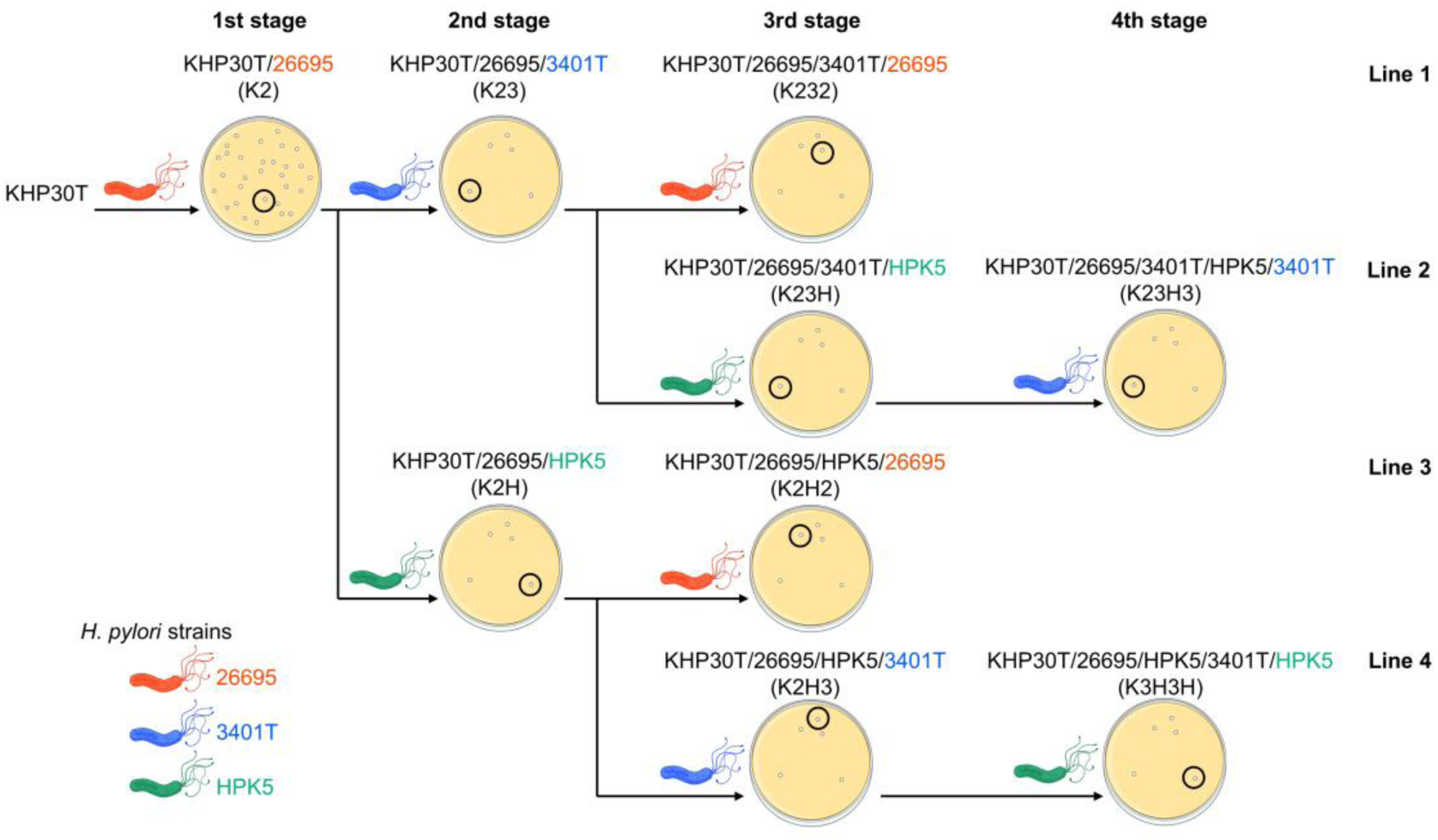
Overview of multistage infection system. Nine KHP30T phages were prepared by adapting to three strains of *H. pylori* (26695, 3401T, HPK5) through four infection stages and four lines. Details of the adaptation procedure are provided in Transparent Methods and Figure S1.

**Figure 2.**
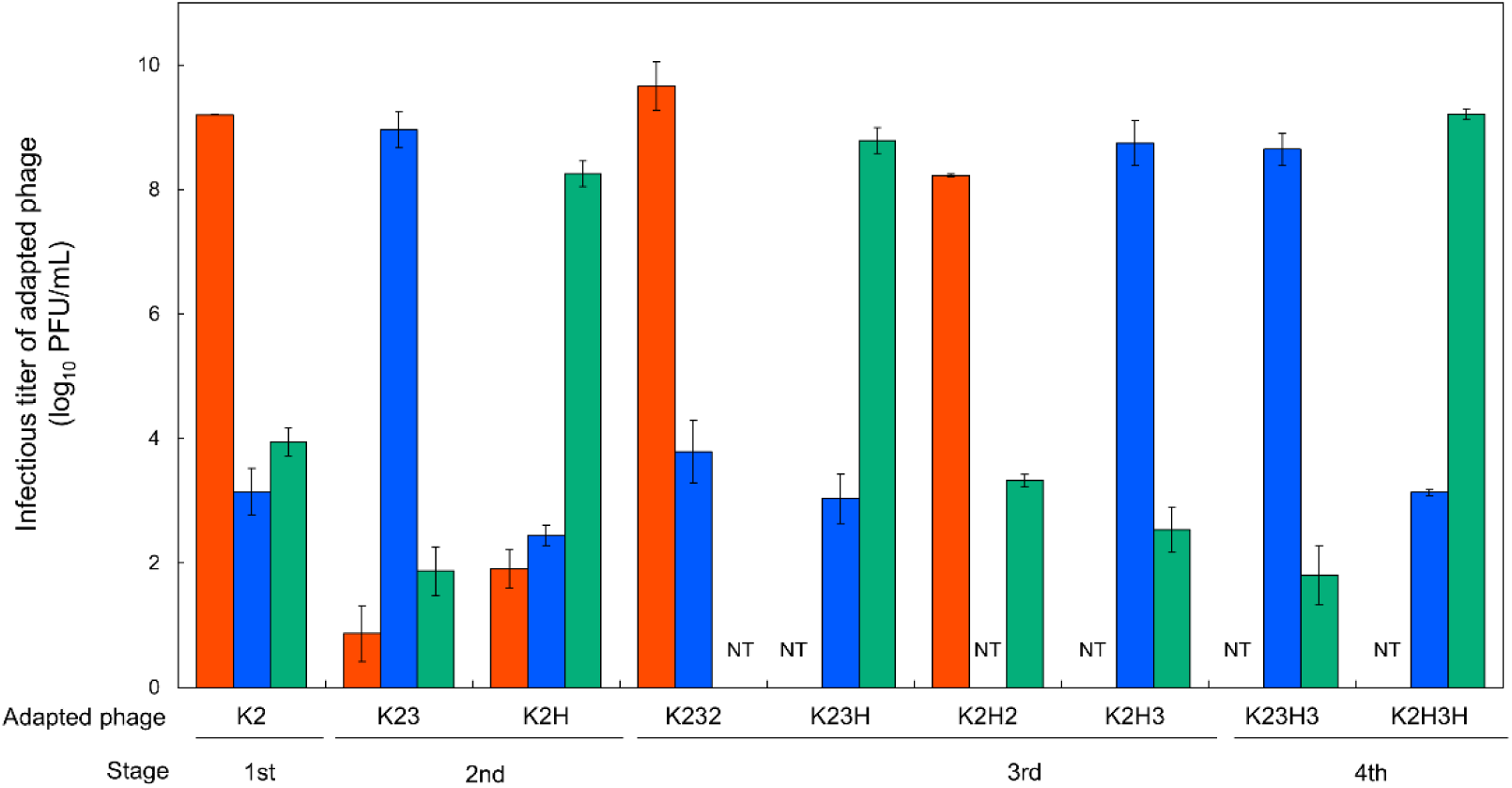
Infectious titer of adapted KHP30T. Orange, blue, and green bars represent KHP30T titers against *H. pylori* 26695, 3401T, and HPK5, respectively. Error bars represent standard deviation (*n* = 3).

Similarly, third-stage adapted phages showed titers against their most recent host strain that were higher than those against other strains. For example, K232 and K2H2 (lines 1 and 3 in Figure 1, respectively) showed significantly higher titers against *H. pylori* 26695 (9.6, and 8.2 log10 PFU/mL, respectively) than those against the other two strains (*p* < 0.01). A similar trend was observed in the fourth stage: K23H3 (line 2) showed a higher titer against 3401T (8.6 log10 PFU/mL) than that against HPK5 (1.8 log10 PFU/mL) (*p* < 0.01). K2H3H (line 4) showed a higher titer against *H. pylori* HPK5 (9.2 log10 PFU/mL) than that against 3401T (3.1 log10 PFU/mL, *p* < 0.01) (Figure 2).

When comparing between adapted phages, the infectious titers against each of the last host strains were generally similar. For example, K2, K232, and K2H2, which all shared *H. pylori* 26695 as their last host, showed titers >8.0 log10 PFU/mL against 26695 (range: 8.2–9.6 log10 PFU/mL) but significantly lower titers against other strains (range: 3.1–3.9 log10 PFU/mL).

Likewise, K23 and K23H3 showed significantly higher titers against 3401T (range: 8.5–8.6 log10 PFU/mL) than those against any other strain. Phages K2H, K23H, and K2H3H also showed significantly higher titers against HPK5 (8.2–9.2 log10 PFU/mL) than those against the others (Figure 2).

Additionally, we measured the infectious titers of the three adapted phages (K2, K23, and K2H) after large-scale cultivation to evaluate whether host tropism changed with passaging. The reproliferated phages exhibited the same trends as the original phages across all host combinations; titers were highest against the most recent host but were lower against the other hosts, including those previously adapted strains (Figure S3). Taken together, these results suggest that the generation gap during reproliferation did not significantly affect the infectivity of the adapted phages in this study.

### Genomic analysis of host H. pylori strains and adapted phages

SMRT sequencing of two *H. pylori* strains (3401T and HPK5) and three adapted phages (K2, K23, and K2H) was performed to determine the epigenomes of both host strains and adapted phages during infection. Genomic and epigenomic data for *H. pylori* 26695 were retrieved from the National Center for Biotechnology Information (NCBI) RefSeq database (accession no. CP003904.1) and a previously published report.^26^ The complete circular genomes of *H. pylori* 3401T and HPK5 were obtained by de novo assembly using high fidelity (HiFi) reads (Table S1). The general characteristics of these genomes (e.g., genome size, coding sequence [CDS] number, and GC content) are similar to those of other *H. pylori* genomes deposited in the NCBI database. The reconstructed *H. pylori* 3401T genome showed >99.99% average nucleotide identity (ANI) with the ancestral strain *H. pylori* 3401 (AP024599.1).^27^ Minor differences in the sequences most likely arose from genomic mutations during serial subculturing and/or from different sequencing technologies and bioinformatic software tools. No extrachromosomal content was observed in any of the strains.

Using functional annotation with REBASE, we predicted 32–36 MTase and 16–21 REase genes in the three *H. pylori* genomes (Tables S1 and S2). The most abundant MTase type was Type II, whereas Types I and III were nearly equivalent. A systematic search for phage defense systems in the genomes revealed 14, 9, and 14 RM systems in *H. pylori* 26695, 3401T, and HPK5, respectively (Table S3). More than half of the detected MTases were orphaned. The most common modification was m6A, followed by m5C and m4C. Although it has been reported that DNA methylation is also involved in mediating other defense systems (e.g., BREX^28^ and DISARM^29^), no other systems associated with DNA modification have been identified. In addition to modification-mediated defense systems, six to nine other systems (e.g., AbiL^30^ and dynamins^31,32^) were identified in the three strains (Table S3).

Mapping KHP30T reads to the original KHP30 phage genome (AB647160.1)^33^ identified an approximately 2-kilobase (kb) insertion in all three phages. No single nucleotide polymorphisms or short insertions/deletions (indels) were detected. The insertion matched a segment of the *H. pylori* 3401 genome, indicating it may have been transferred from the host during serial passaging. This insertion likely does not significantly affect the general characteristics of the phage, such as infection efficiency or host tropism, consistent with the titration results above (Figure 2) and previous reports.^33–35^ The complete circular genome of KHP30T was reconstructed using HiFi reads from the K2 phages (Table S1). Read mapping analysis showed no obvious genomic variations fixed within the phage population. No MTase or REase genes, as well as phage defense systems, were detected in the phage genome.

### Epigenomic analysis of H. pylori strains and adapted phages

We successfully identified numerous base modifications across both the *H. pylori* and phage genomes (Figures 3 and S4, Table S4). From motif predictions, we detected 15 and 20 motifs in the *H. pylori* 3401T and HPK5 genomes, respectively, as well as 17 motifs in *H. pylori* 26695, the latter retrieved from a previous study^26^ (Table S4). Among these motifs, the most abundant modification type was m6A, followed by m4C and m5C. There was a slight discrepancy in the modification types of the encoded MTases, possibly due to the SMRT sequencing’s low detection sensitivity for m5C modifications. As expected, the detected m5C motifs showed low modification ratios (<50.0%) across all three *H. pylori* strains. Only two motifs, C**A**<u>T</u>G and G**A**<u>T</u>C (with the modified bases in bold and their reverse complements underlined), were shared among all three strains. However, 28 of 39 (72%) motifs were present in either of the 3 strains, indicating high diversity in methylation patterns across these strains. This trend aligns with previous findings that *H. pylori* strains possess highly divergent combinations of modified motifs.^26,36^

**Figure 3.**
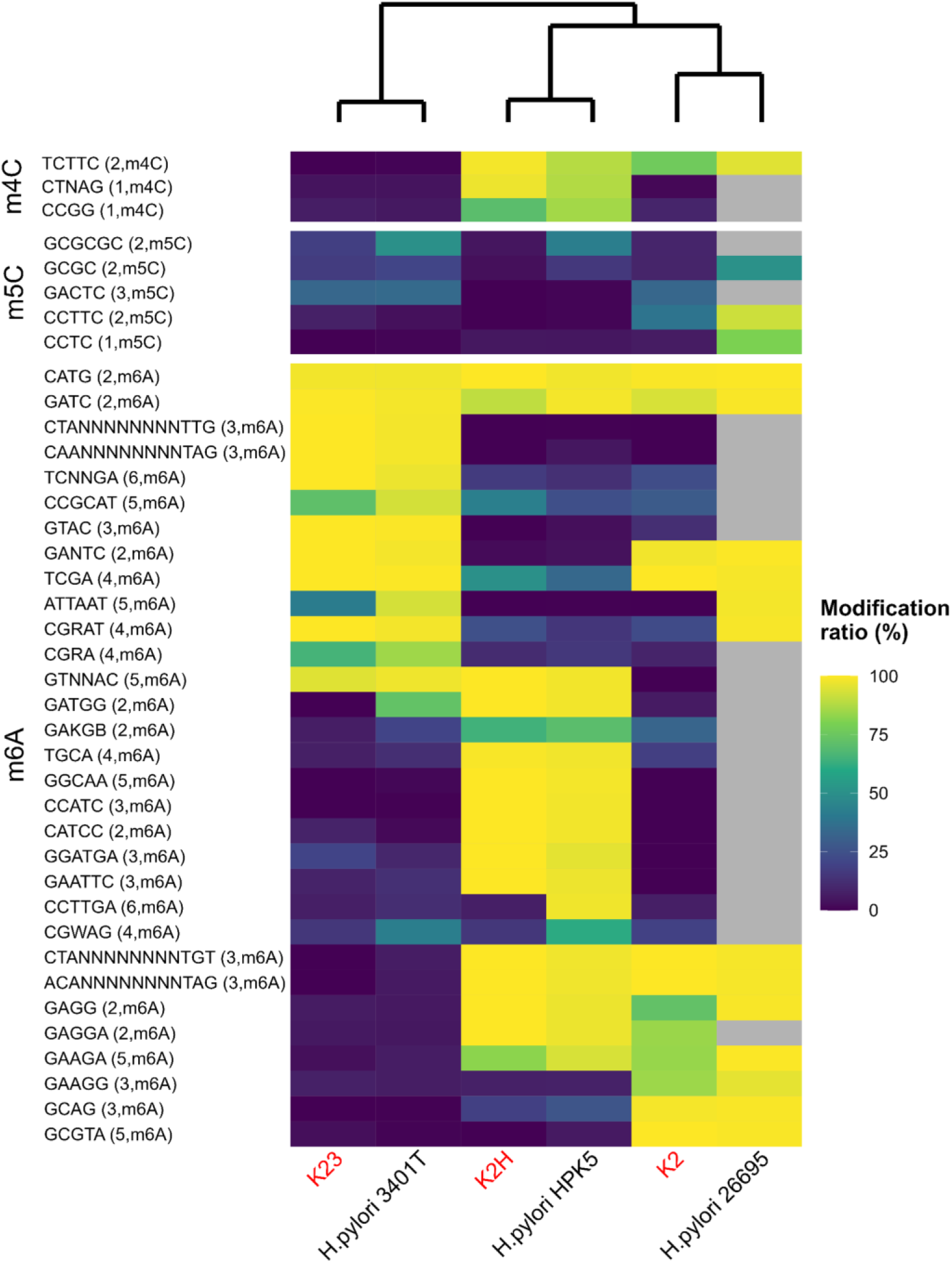
Methylomes of adapted phages and *H. pylori* strains. Methylated motifs were predicted from the three *H. pylori* strains. Motifs are shown separately according to methylation type. Phages and *H. pylori* strains are colored in red and black, respectively, in the bottom text. Text on the left side indicates the methylated motifs with methylated nucleotide position and methylation type. The top dendrogram was calculated using Euclidean distance and Ward method. For *H. pylori* 26695, methylated motifs and ratios were retrieved from the previous study reported by Krebes et al. (2014), and gray cells indicate motifs for which no data are available. Some motifs showed comparatively high modification ratios due to motif overlap. For example, the modification ratio of CGR**A**T (R=A/G) in *H. pylori* 3401T is explained by motif overlapping with CGR**A**, which was detected from epigenomic analysis of the strain. In addition, CGW**A**G (W=A/T) in *H. pylori* 3401T partly overlapped with CGR**A**, i.e., CGA**A**G. A motif G**A**GG in *H. pylori* HPK5 also overlapped with the detected G**A**GGA from the strain, and in contrast, G**A**GGA in K2 overlap with G**A**GG reported from *H. pylori* 26695 previously. Among all seven CCGC**A**T sites in K23, two and one overlapped with other motifs C**A**<u>T</u>G (i.e., CCGC**A**TG) and **A**TCC (i.e., CCGC**A**TCC), respectively. Hence, CCGC**A**T showed a comparatively higher modification ratio in K23.

Next, we compared the methylation patterns in *H. pylori* strains and their adapted phages. Overall, the methylated motifs were shared between pairs of *H. pylori* strains and their respective adapated phages (Figure 3). For example, among the motifs detected in *H. pylori* HPK5, with the exception of m5C motifs, 17 of 19 were highly modified in phage K2H (motif modification ratios >68.9% and >63.6% in the host and phage genomes, respectively). Similarly, 12 of 13 motifs were shared between *H. pylori* 3401T and phage K23 with ratios >73.9% and >41.7%, respectively. In addition, 12 of 14 motifs were concordantly methylated between *H. pylori* 26695 and phage K2, with ratios >95.2% and >71.3%, respectively. In contrast, sets of methylated motifs were not shared between non-corresponding host-phage pairs.

Notably, several motifs were identified that were highly methylated in the host genomes but showed minimal methylation in the adapted phage genomes (Figure 3). For instance, the m6A-type motif G**A**TGG was methylated in the *H. pylori* 3401T genome with a high modification ratio (73.9%), but was undetected in the K23 genome. Similarly, the motif CCTTG**A** was highly modified in *H. pylori* HPK5 (81.1%) but not in K2H (7.1%). Additionally, motifs A<u>T</u>TA**A**T and CGR**A**T have been reported to be methylated at high modification ratios (98.5% and 98.9%, respectively) in *H. pylori* 26695^26^ but showed much lower modification in phage K2 (0.0% and 22.6%, respectively).

### Phase-variable RM systems and associated genes

In the *H. pylori* HPK5 genome, a G9-tract was identified within the HPYHPK5_00170−00190 region (Figure S5A), displaying high sequence similarity to the M.Hpy300X Type III m6A MTase gene. This gene is known to induce methylome variation through simple sequence repeats (SSRs).^37^ The basal state of the poly-G tract results in a stop codon and a truncated protein product. However, the addition of a single nucleotide to the tract results in a frameshift, generating a fully functional fusion protein. Similar to the Hpy300X system, this tract likely modulates MTase activity, suggesting that HPYHPK5_00150-00160 functions as a phase-variable Type III RM system in conjunction with a Type III REase (HPYHPK5_00170). Similar structures were also predicted in *H. pylori* 26695, specifically in Type III RM systems AFV42588−42590 (HpyAXVII) (Figure S5B) and AFV42742−42744 (Figure S5C), with the former previously characterized.^38^

In the *H. pylori* 3401T genome, HPY3401T_09270 and HPY3401T_09280 harbor the C19- and C18-tracts, respectively (Figure S5D). These sequences showed the highest similarity to the sequences of genes associated with the HpyAXVI system (89.8% and 89.0% identity to HpyAXVI-mut2 and HpyAXVI-mut1, respectively). The HpyAXVI system, composed of Type II RM genes (REase and MTase fusion genes), was originally identified in *H. pylori* 26695^26^ (Figure S5E). This system is inactive when a stop codon separates the two CDSs fillowing the first tract but becomes active when a frameshift near the tract fuses the two CDSs. When active, the tract produces a truncated protein that regulates MTase sequence specificity. Therefore, HPY3401T_09270 and HPY3401T_09280 are likely phase-variable Type II RM systems. In addition, HPYHPK5_13900 and HPYHPK5_13910 contain C14- and G15-tracts, respectively (Figure S5F). The former showed the highest sequence similarity (88.2%) to Hpy300XI found in *H. pylori* BCM-300 strain. This system uses two tracts to facilitate phase variation, likely operating similarly to HpyAXVI^26^. Thus, HPYHPK5_13900 and HPYHPK5_13910 may also comprise phase-variable Type II RM systems.

Beyond MTases, certain SSRs were associated with REase and specificity (S) subunit genes. In all *H. pylori* genomes, four tracts were identified within Type I REase CDSs (Figure S5G, S5H, S5I, and S5J). In contrast, cognate MTases and S genes were not associated with SSRs. As Type I MTases require an S subunit complex for functionality, these tracts likely impact only the cleavage activity of Type I RM systems, leaving methylation functions unaffected. In addition, a C17-tract was found in the HPY3401T_11810 REase, part of a Type III RM system with the HPY3401T_11820 MTase (Figure S5K). While Type III MTases function independently, REases require a cognate MTase for cleavage. Tandem short-sequence repeats, including SSRs, are common in Type III MTase genes and act as ON/OFF switches regulating phase variation,^39^ as also observed in this study (Figure S5A, S5B, and S5C). Therefore, this C17-tract may specifically regulate only REase activity, while the cognate MTase may be constituitively active. AFV41850 from *H. pylori* 26695 and HPY3401T_02460 from 3401T showed the highest sequence similarity to Type IV REase EcoCTGmrSD and possessed T15- and T19-tracts located 47 and 46 base paris (bp) upstream of their CDSs, respectively (Figure S5L and S5M). Type IV REases cleave methylated DNA strands in contrast to other REase types, suggesting that these tracts likely regulate the ON/OFF switch for Type IV RM systems. Finally, consistent with a previous study’s findings, we confirmed that G14-tracts within the S gene comprise a BcgI-like Hpy99XXII system accompanied by an upward-adjusted RM gene, though its function is still unclear^26^ (Figure S5N).

## DISCUSSION

DNA methylation-mediated biological processes affect a wide range of microbial ecologies, including phage-host interactions. There is growing interest in epigenomic systems of diverse prokaryotes and viruses, given their importance in microbial physiology, genetics, evolution, and ecology.^2,40,41^ To investigate the role of methylation in phage-host interactions, *H. pylori* and phage KHP30 were chosen as model organisms.

*H. pylori* is a major human pathogen that causes chronic inflammation in the stomach and gastric cancer. It harbors multiple RM systems with highly variable composition, resulting in diverse methylomes across strains.^26,36,42–44^ However, epigenomic evaluation has not been performed to determine whether the MTases encoded in the *H. pylori* genome play a role against invading phages. We previously reported that KHP30, a double-stranded DNA phage belonging to the *Schmidvirus* genus, infects *H. pylori*^33–35^ and has potential as a therapeutic agent against antibiotic-resistant *H. pylori*, owing to its broad host range.^34^ However, a significant barrier to phage therapy for *H. pylori* lies in the limited understanding of its defense mechanisms against KHP30 infection.

To clarify the relationship between DNA methylation and phage-host interactions, we designed a multistage infection system (Figure 1) and analyzed the methylomes of three *H. pylori* strains (26695, 3401T, and HPK5) along with three adapted KHP30T phages (K2, K23, and K2H) with varying infection histories. The adapted phages showed high titers against their most recent host strain but significantly lower titers against other strains, even if they had previously adapted to them (Figure 2). Considering the short generation time during cultivation, gene expression disruptions or enzymatic inactivation from mutations were unlikely to affect tropism significantly. As expected, the reproliferated phages consistently showed high titers only against the latest infected host (Figure S3).

Our epigenomic analysis revealed that each *H. pylori* strain contained distinct sets of MTase genes, many positioned near REase genes within the RM systems (Tables S1 and S2). Adapted phages were found to share methylated motifs with their final adapted host strain, indicating that phage epigenomes were modified by host MTases during infection. Taken together, our work demonstrates that adapted phages acquire DNA methylation from their host to evade from the host’s RM systems, thereby increasing infection efficiency, as has been previously hypothesized.^45^ To the best of our knowledge, this is the first study to directly observe epigenomic changes in the phage genome during infection (Figure S4). Although methylated motifs were shared between adapted phages and their host, exceptions were noted. Specifically, several motifs were methylated in the host genome but not in the adapted phage (Figure 3). These exceptions may partially be due to phase variations in several RM systems of *H. pylori* (Figure S5).

Phase variation is a random and reversible switching mechanism in gene expression that generates transcriptional, translational, and/or phenotypic diversity within bacterial populations.^46^ Bacterial MTases within RM systems often undergo phase variation, resulting in heterogeneous cells with varied RM activity within a clonal population.^47^ This heterogeneity enables phages to selectively infect cells with reduced RM activity, as their epigenomes are methylated by the MTases in these hosts. A primary mechanism regulating phase variation is the presence of SSRs within or upstream of RM genes, with phase-variable ON/OFF switching occuring in the former, resulting in reversible changes in the number of SSRs.^47–51^ Our analysis found several phase-variable regulons or phasevarions associated with RM genes in all three *H. pylori* genomes (Figure S5). Further single-cell evaluation such as combining immunofluorescence imaging to track phase-variated proteins^52^ with fluorescence *in situ* hybridization for phage detection, are necessary to clarify the link between phase variation and phage variation. Moreover, bioinformatic tools for detecting modifications needs to be improved, as current modification detection tools do not account for phase-variable epigenomes and instead treat modifications on a binary, population-wide manner. Another possible explanation for the methylated motifs found exclusively in the host genome is the potential presence of anti-RM systems in the phage that block host MTase activity. For example, DarA and DarB in P1 phages protect recognition sites on their genome from host RM activity^53,54^. Similarly, Ocr in T7 phages mimics DNA structure to block the DNA-binding groove of Type I REases/MTases.^55^ However, no genes associated with known anti-RM systems^56^ were found in the KHP30T genome.

Defense systems are involved in the evolutionary arms race between phages and hosts.^57^ Epigenomic modifications significantly influence their co-evolution and promote microbial diversification.^58,59^ Our findings revealed that the phage epigenome was altered by host bacteria during infection, enhancing the phage’s ability to reinfect the same host strain (Figures 2 and 3). To counter the epigenomic synchronization of invading phages and avoid extinction of the entire population by phage persistence, some bacteria employ variable phage-defense systems. For example, many species of gut bacteria, such as *Bacteroides fragilis* and *H. pylori*, show strain-specific variations in gene sets associated with phage defense, including MTase genes.^60–62^ While gene acquisition or deletion rely on genomic events such as horizontal gene transfer and recombination, such events occur at infrequently. Alternatively, a more dynamic strategy by bacteria involves phase variation to prevent phage circulation. For example, *Vibrio cholerae* modulates the expression of the lipopolysaccharide O1 antigen through phase variation to escape predation by O1-specific phages.^63^ In *H. pylori*, inter-strain variation in methylomes caused by MTase phase variation has been reported.^36,37^ Therefore, antagonistic coevolution between prokaryotes and phages, along with associated epigenetic changes, is of great interest for understanding the diversity within microbial ecosystems.

In summary, our study demonstrates that phages can overcome host RM systems by acquiring methylation modifications from host-encoded MTases during infection. Adapted phages showed significantly higher titers against their most recent host strain, likely due to neutralization of the host RM system through acquired methylation. Our results enhance our understanding of DNA methylation-mediated phage–host interactions and their impact on microbial ecology across diverse environments.

Our approach provides new insights into the design of therapeutic phage cocktails. Phage therapy holds promise across various fields, including human and plant diseases,^64–67^ owing to phages’ advantages such as host specificity, self-amplification, biofilm degradation, and low toxicity to humans.^68,69^ One of the primary challengess in phage therapy is bacterial resistance; however, phage cocktails, as shown in studies on *E. coli* O157:H7 and *Klebsiella pneumoniae*, offer a strategy to combat resistance by incorporating diverse phage variants.^70,71^ We propose that a cocktail of adapted phages, each with distinct methylation motifs, could improve the chances of overcoming host modification-mediated defense systems, such as RM systems. Further research to understand the mechanism of phage adaptation and evaluate their therapeutic potential will contribute to the development of tailored phage cocktails for medical and industrial applications.

### Limitation of the study

This study has few limitations. First, we did not experimentally verify the activity and specificity of the MTases and REases identified, leaving uncertainty about which RM systems are actively involved in phage defense and the specific MTases responsible for the modified motifs. Second, although SMRT sequencing technology efficiently identifies m6A and m4C modifications,^26^ it has limited sensitivity for detecting m5C modifications, one of the most common methylation types in nature after m6A and m4C. In total, five m5C motifs were detected across all three *H. pylori* strains (Table S4), but they showed low modification ratios (Figure 3, Table S4), presumably due to the low detection accuracy for this modification type using current SMRT technology. In addition to methylation, other epigenetic modifications, such as phosphorothioate modification, have recently been linked to phage defense.^72^ Expanding sequencing technologies to detect a wider range of DNA modifications is crucial for a comprehensive understanding of epigonomic functions. Third, although HiFi reads are highly accurate (>99.9%), sequencing of homopolymer regions remains challenging,^73–75^ complicating estimatations of tract diversity. Nonetheless, given the high genome coverage achieved in this study (Table S1), the tracts associated with RM genes (Figure S5) likely represent the predominant phase variation state in the populations studied.

## RESOURCE AVAILABILITY

### Lead contact

Further information and requests for resources and reagents should be directed to and will be fulfilled by the lead contact, Michiko Takahashi.

### Data availability

The assembled genomes were deposited in DDBJ/ENA/GenBank (S1 Table). All data are registered under BioProject ID PRJDB17787 [https://ddbj.nig.ac.jp/resource/bioproject/PRJDB17787]. Any additional information required to reanalyze the data reported in this paper is available from the lead contact upon request.

## METHODS

All methods can be found in the Transparent Methods.

## ACKNOWLEDGMENTS

The authors are greatful to Waka Ishida, Yusaku Funaoka, and Takahiro Ishikawa for experimental support, and to Daichi Morimoto for helpful comments on this work. This study was supported by JSPS KAKENHI (grant numbers JP20K15582 and JP22K08875), ACT-X, Japan Science and Technology Agency (grant numbers JPMJAX21BD and JPMJAX22BK), and the budget of Kochi University Hospital (Japan). The experiments were conducted at the Division of Biological Research, Science Research Center, Kochi University. Computations were partially performed on the supercomputer system at the Institute for Chemical Research, Kyoto University, and the NIG supercomputer system at ROIS National Institute of Genetics, and the Data Analysis and the Earth Simulator systems at JAMSTEC.

## AUTHOR CONTRIBUTIONS

MT designed the study, acquired funding, performed the experiments, and drafted the manuscript. SH designed the study, performed the bioinformatics analyses, and wrote the manuscript. YM, RS, and TU performed the experiments. HM acquired funding and provided resources. SS, KN, and HT supervised the project. SM conceived the original idea, acquired funding, and supervised the project. All authors have read and approved the final manuscript.

## SUPPLEMENTAL INFORMATION

**Figure S1. Overview of preparation procedure for adapted phages.**

**Figure S2. Tukey-Kramer pairwise comparisons of phage titers.** Corresponding titers are shown in Figure 2. Combination of phage and host strain are indicated as ‘adapted phage’: ‘host strain used for titration’ format in the x-axis: for example, ‘K2:2’ means a titer of K2 against *H. pylori* 26695. Solid black bar within each box represents median. Upper alphabet summarizes the results of the Tukey-Kramer test; letters denote significant differences (*p* < 0.01) in mean titers with groups a > b > c > d > e > f.

**Figure S3. Infectious titer of reproliferated phages.** (A) Orange, blue, and green bars represent titer of reproliferated KHP30T phages to *H. pylori* 26695, 3401T, and HPK5, respectively. Error bars indicate standard deviation (*n* = 3). (B) Tukey-Kramer pairwise comparisons of reproliferated phage titers. Phage and host combinations are indicated in the same format as Figure S2. Infectious titer of the ‘Original phages’ is derived from Figure 2. Solid black bar within each box represents median. Upper alphabet summarizes the results of the Tukey-Kramer test; letters denote significant differences (*p* < 0.01) in mean titers with groups a > b > c > d > e > f.

**Figure S4. Methylomes of adapted phages.** Circos plot showing the distribution of DNA methylation. Orange, blue, and red bars represent m4C, m5C, and m6A modifications, respectively. K2 and K23 were sequenced using PacBio Sequel II in CCS mode, whereas K2H was sequenced using PacBio Sequel in CLR mode.

**Figure S5. Genetic organization of RM system genes associated with SSRs.** Homopolymeric tracts inside or upstream of RM genes were specified as SSRs. RM gene types are indicated by color and text inside to each CDS. CDSs not assigned to RM system genes are colored in gray.

**Table S1. Descriptions and statistics of *H. pylori* strains and KHP30T phages.**

**Table S2. Detected RM system genes in *H. pylori* strains.**

**Table S3. Predicted phage defense systems in *H. pylori* strains.**

**Table S4. Methylated motifs predicted from *H. pylori* strains.**

**Transparent Methods**

