## Supplemental information for "Host-encoded DNA methyltransferases modify the epigenome and host tropism of invading phages"

### Supplemental Figures

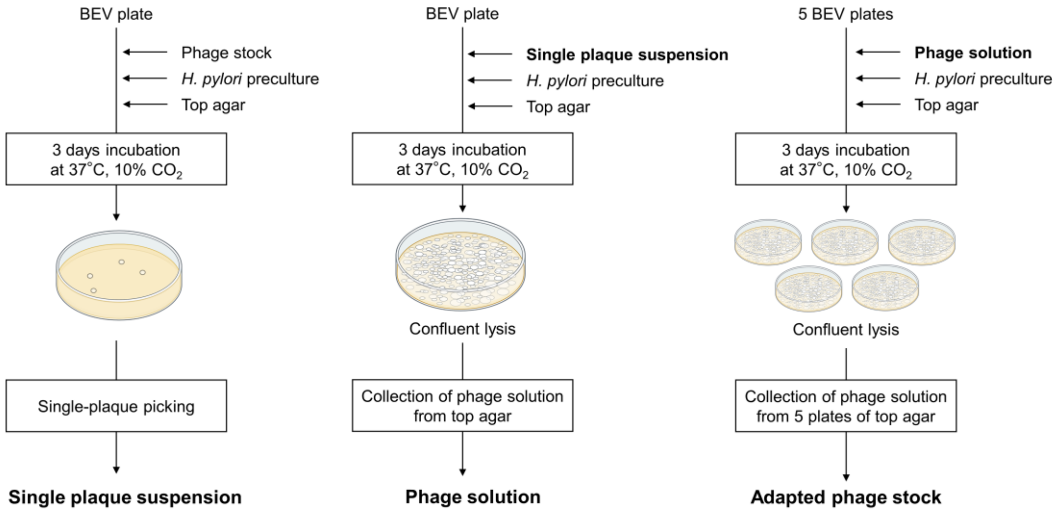


**Figure S1. Overview of preparation procedure for adapted phages.**

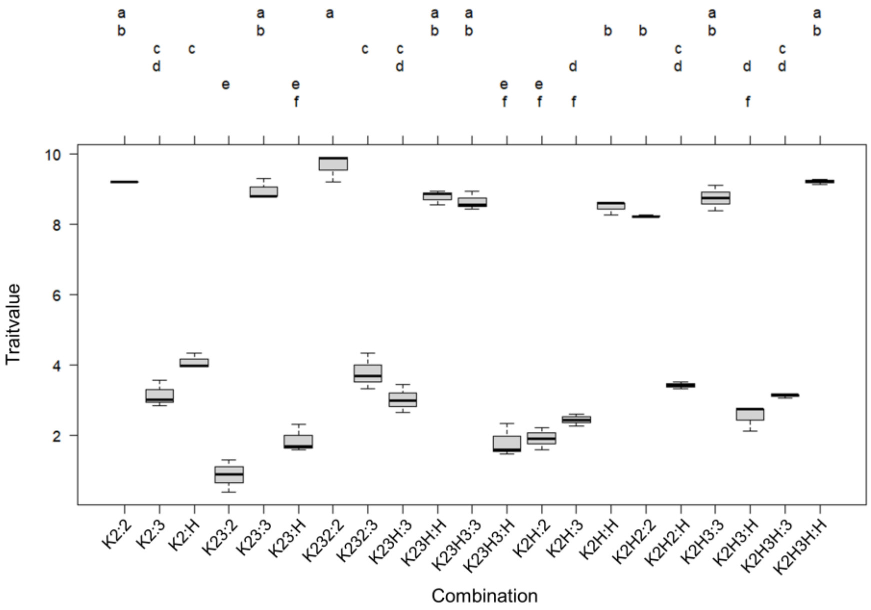


**Figure S2. Tukey-Kramer pairwise comparisons of phage titers.** Corresponding titers are shown in Figure 2. Combination of phage and host strain are indicated as 'adapted phage': 'host strain used for titration' format in the x-axis: for example, 'K2:2' means a titer of K2 against *H. pylori* 26695. Solid black bar within each box represents median. Upper alphabet summarizes the results of the Tukey-Kramer test; letters denote significant differences ($p$ < 0.01) in mean titers with groups a > b > c > d > e > f.

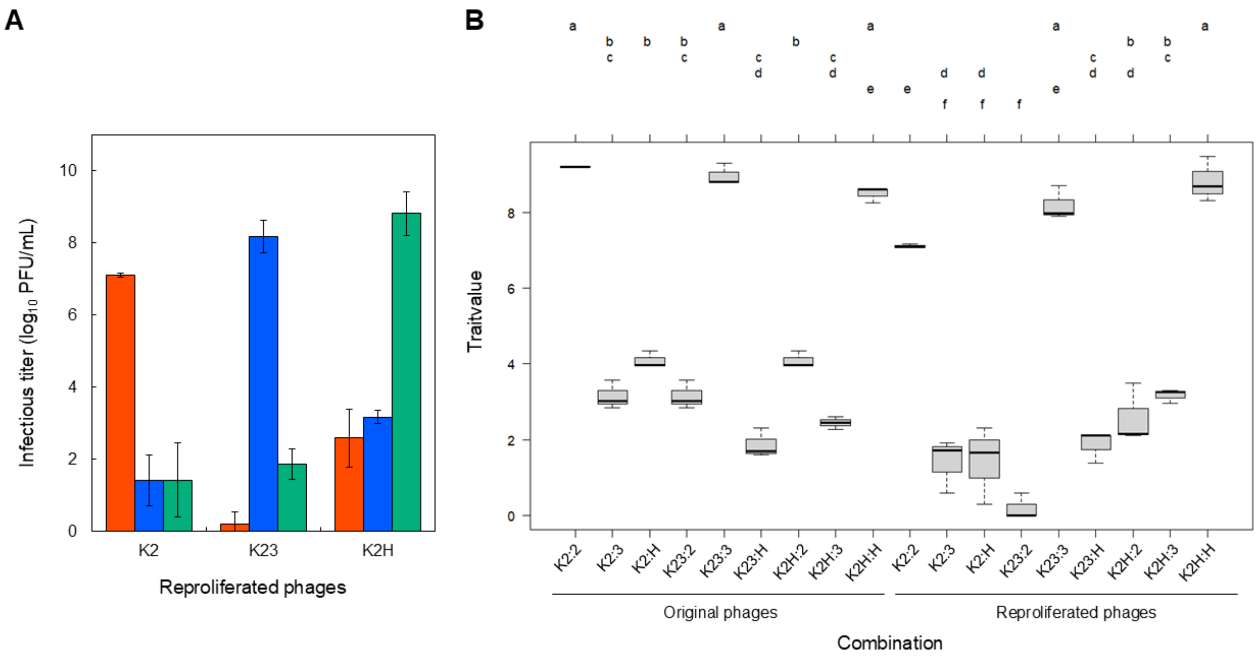


**Figure S3. Infectious titer of reproliferated phages.** (A) Orange, blue, and green bars represent titer of reproliferated KHP30T phages to *H. pylori* 26695, 3401T, and HPK5, respectively. Error bars indicate standard deviation ($n$ = 3). (B) Tukey-Kramer pairwise comparisons of reproliferated phage titers. Phage and host combinations are indicated in the same format as Figure S2. Infectious titer of the 'Original phages' is derived from Figure 2. Solid black bar within each box represents median. Upper alphabet summarizes the results of the Tukey-Kramer test; letters denote significant differences ($p < 0.01$) in mean titers with groups a > b > c > d > e > f.

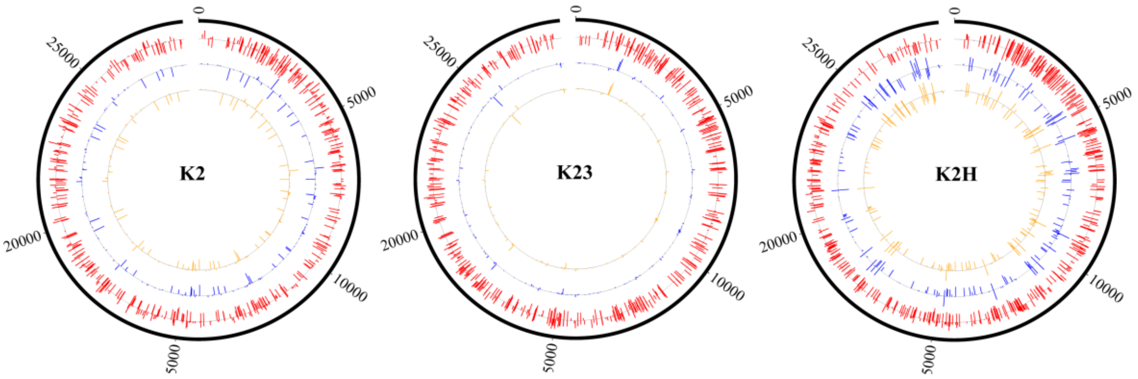


**Figure S4. Methylomes of adapted phages.** Circos plot showing the distribution of DNA methylation. Orange, blue, and red bars represent m4C, m5C, and m6A modifications, respectively. K2 and K23 were sequenced using PacBio Sequel II in CCS mode, whereas K2H was sequenced using PacBio Sequel in CLR mode.

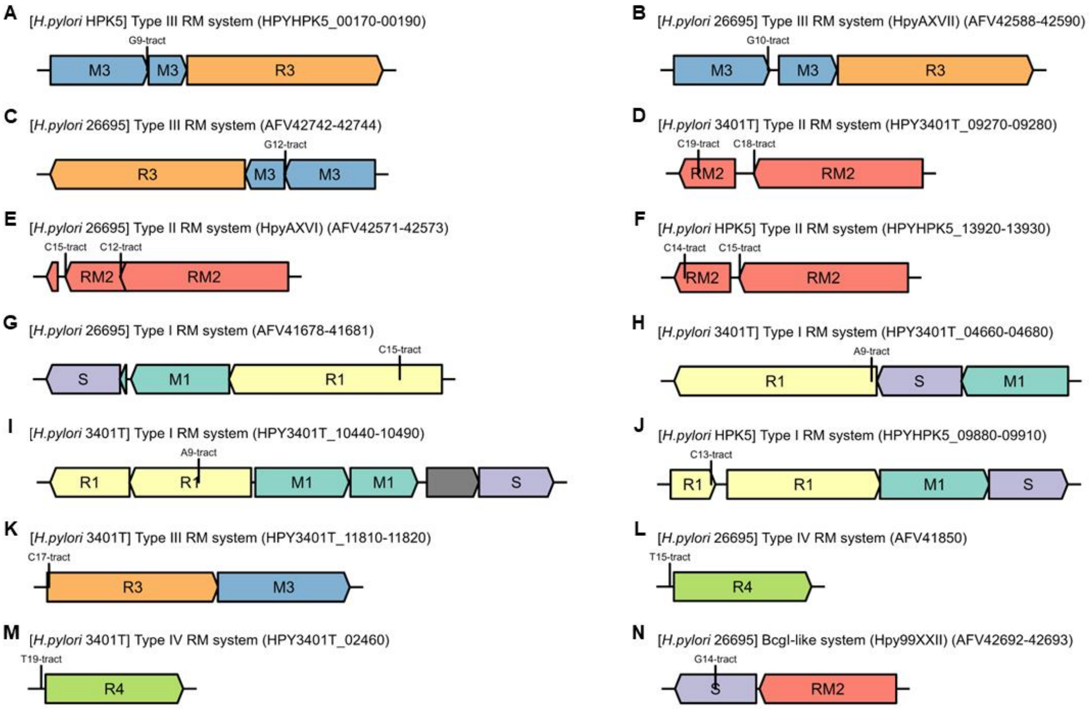


**Figure S5. Genetic organization of RM system genes associated with SSRs.** Homopolymeric tracts inside or upstream of RM genes were specified as SSRs. RM gene types are indicated by color and text inside to each CDS. CDSs not assigned to RM system genes are colored in gray.

#### Supplemental Tables

**Table S1. Descriptions and statistics of *H. pylori* strains and KHP30T phages.**

| Organism | *Helicobacter pylori* 26695 | *Helicobacter pylori* 3401T | *Helicobacter pylori* HPK5 | Helicobacter phage KHP30T (K2) | Helicobacter phage KHP30T (K23) | Helicobacter phage KHP30T (K2H) |
|---|---|---|---|---|---|---|
| Platform | ND | PacBio Sequel II | PacBio Sequel II | PacBio Sequel II | PacBio Sequel II | PacBio Sequel |
| Sequencing mode | ND | HiFi | HiFi | CCS | CCS | CLR |
| Subreads | ND | ND | ND | 450,651 | 1,265,358 | 60,610 |
| ---Read length (bp) | ND | ND | ND | 4,892 ± 4,443 | 4,065 ± 3,111 | 5439 ± 2759 |
| ---Total base (bp) | ND | ND | ND | 2,204,427,929 | 5,143,821,232 | 329,645,020 |
| HiFi reads | ND | 84,112 | 104,380 | 37,903 | 95,126 | ND |
| ---Read length (bp) | ND | 4,926 ± 771 | 4,311 ± 540 | 7,186 ± 6,278 | 5,335 ± 3,674 | ND |
| ---Total base (bp) | ND | 801,415,974 | 1,068,931,514 | 272,376,564 | 507,522,297 | ND |
| Assembled genome | 1 | 1 | 1 | 1 | ND | ND |
| ---Genome size (bp) | 1,667,892 | 1,591,257 | 1,583,340 | 28,242 | ND | ND |
| ---Coverage (subreads) | ND | ND | ND | ND | ND | 10,608.6 |
| ---Coverage (HiFi reads) | ND | 1,005.4 | 1,328.1 | 6,638.1 | 10,452.0 | ND |
| ---GC content (%) | 39 | 38.79 | 38.79 | 36 | ND | ND |
| ---CDSs | 1,577 | 1,538 | 1,514 | 31 | ND | ND |
| --- ---MTase | 36 | 32 | 34 | 0 | ND | ND |
| --- ---REase | 16 | 21 | 21 | 0 | ND | ND |
| ---RM system | 14 | 10 | 14 | 0 | ND | ND |
| ---modified motifs | 17 | 15 | 20 | ND | ND | ND |
| ---Accession number | CP003904.1 | AP031459.1 | AP031460.1 | LC813552.1 | ND | ND |
| Reference of genome | NA | This study | This study | This study | This study | This study |

ND: no data. NA: Not available.

#### Table S2. Detected RM system genes in *H. pylori* strains.

| Strain | CDS ID | SSR | SSR start | SSR end | Top-hit protein in REBASE | Identity (%) | RM type | Modification type | Recognition motif of the closest-match MTase | Modification position of the motif |
|---|---|---|---|---|---|---|---|---|---|---|
| *Helicobacter pylori* 26695 | NC_018939.1_46 | | | | M1.HpyAVI | 100.0 | M2 | m6A | CCTC | -2 |
| *Helicobacter pylori* 26695 | NC_018939.1_47 | | | | M.HpyD27I | 99.4 | M2 | m5C | CCTC | 1 |
| *Helicobacter pylori* 26695 | NC_018939.1_50 | | | | HpyAV | 100.0 | R2 | NA | CCTTC | NA |
| *Helicobacter pylori* 26695 | NC_018939.1_51 | | | | M.HpyAV | 100.0 | M2 | m5C | CCTTC | 2 |
| *Helicobacter pylori* 26695 | NC_018939.1_89 | | | | HpyAIII | 100.0 | R2 | NA | GATC | NA |
| *Helicobacter pylori* 26695 | NC_018939.1_90 | | | | M.HpyAIII | 100.0 | M2 | m6A | GATC | 2 |
| *Helicobacter pylori* 26695 | NC_018939.1_263 | | | | M.HpyAX | 99.7 | M2 | m6A | TCGA | 4 |
| *Helicobacter pylori* 26695 | NC_018939.1_266 | | | | M.Hpy66IX | 97.2 | M2 | Nm4C | CCGG | 1 |
| *Helicobacter pylori* 26695 | NC_018939.1_367 | | | | HpyGI | 97.0 | R2 | NA | TCNNGA | NA |
| *Helicobacter pylori* 26695 | NC_018939.1_368 | | | | M.HpyGI | 98.2 | M2 | NA | TCNNGA | NA |
| *Helicobacter pylori* 26695 | NC_018939.1_369 | | | | M.HpyGI | 98.3 | M2 | NA | TCNNGA | NA |
| *Helicobacter pylori* 26695 | NC_018939.1_457 | | | | S.HpyUM032XVI | 92.9 | S | NA | CCANNNNNNTC | NA |
| *Helicobacter pylori* 26695 | NC_018939.1_458 | | | | M.Hpy66XV | 96.9 | M1 | m6A | CAGNNNNNTC | 2 |
| *Helicobacter pylori* 26695 | NC_018939.1_459 | | | | M.HpyUM032XVI | 92.6 | M1 | m6A | CCANNNNNNTC | 3 |
| *Helicobacter pylori* 26695 | NC_018939.1_460 | C15 | 485322 | 485336 | DvuI | 27.1 | R1 | NA | TGACNNNNNNTTC | NA |
| *Helicobacter pylori* 26695 | NC_018939.1_474 | | | | M.HpyAVII | 100.0 | M2 | m6A | ATTAAT | 5 |
| *Helicobacter pylori* 26695 | NC_018939.1_477 | | | | M.Hpy66II | 96.2 | M2 | Nm4C | ACNGT | 2 |
| *Helicobacter pylori* 26695 | NC_018939.1_478 | | | | M.Hpy66II | 98.8 | M2 | Nm4C | ACNGT | 2 |
| *Helicobacter pylori* 26695 | NC_018939.1_480 | | | | M.Hpy99XI | 95.7 | M2 | m5C | ACGT | 2 |
| *Helicobacter pylori* 26695 | NC_018939.1_481 | | | | M.Hpy99XI | 95.4 | M2 | m5C | ACGT | 2 |
| *Helicobacter pylori* 26695 | NC_018939.1_510 | | | | EcoCTGmrSD | 25.8 | R4 | NA | NA | NA |
| *Helicobacter pylori* 26695 | NC_018939.1_591 | | | | NgoAX | 28.6 | R3 | NA | CCACC | NA |
| *Helicobacter pylori* 26695 | NC_018939.1_592 | | | | M.HpyAXI | 100.0 | M3 | m6A | GCAG | 3 |
| *Helicobacter pylori* 26695 | NC_018939.1_629 | T15 (47 bp upstream) | 675073 | 675087 | EcoCTGmrSD | 23.6 | R4 | NA | NA | NA |
| *Helicobacter pylori* 26695 | NC_018939.1_668 | | | | HpyAXVIIIC | 100.0 | RM2 | NA | GGANNAG | NA |
| *Helicobacter pylori* 26695 | NC_018939.1_780 | | | | S.HpyAXIII | 100.0 | S | NA | CTANNNNNNNNTGT | NA |
| *Helicobacter pylori* 26695 | NC_018939.1_836 | | | | Hpy99XVI | 90.9 | R1 | NA | RTAYNNNNNRTAY | NA |
| *Helicobacter pylori* 26695 | NC_018939.1_838 | | | | S.Hpy99XVI | 94.4 | S | NA | RTAYNNNNNRTAY | NA |
| *Helicobacter pylori* 26695 | NC_018939.1_839 | | | | S.HpyAXIII | 61.1 | S | NA | CTANNNNNNNNTGT | NA |
| *Helicobacter pylori* 26695 | NC_018939.1_840 | | | | M.HpyAXIII | 100.0 | M1 | m6A | CTANNNNNNNNTGT | 3 |
| *Helicobacter pylori* 26695 | NC_018939.1_901 | | | | M.HpyUM032XI | 96.1 | M2 | m6A | GTNNAC | 5 |
| *Helicobacter pylori* 26695 | NC_018939.1_975 | | | | M.Hpy188II | 48.2 | M2 | m6A | CATG | 2 |
| *Helicobacter pylori* 26695 | NC_018939.1_979 | | | | HpyGI | 83.3 | R2 | NA | TCNNGA | NA |
| *Helicobacter pylori* 26695 | NC_018939.1_1113 | | | | M.HpyAVIII | 100.0 | M2 | m5C | GCGC | 2 |
| *Helicobacter pylori* 26695 | NC_018939.1_1205 | | | | M.HpyAI | 100.0 | M2 | m6A | CATG | 2 |
| *Helicobacter pylori* 26695 | NC_018939.1_1348 | | | | HpyAIV | 100.0 | R2 | NA | GANTC | NA |
| *Helicobacter pylori* 26695 | NC_018939.1_1349 | | | | M.HpyAIV | 100.0 | M2 | m6A | GANTC | 2 |
| *Helicobacter pylori* 26695 | NC_018939.1_1350 | | | | HpyAXVI-mut2 | 100.0 | RM2 | m6A | CRTCNA | 6 |
| *Helicobacter pylori* 26695 | NC_018939.1_1351 | C15 | 1413744 | 1413758 | HpyAXVI-mut1 | 100.0 | RM2 | m6A | CRTTAA | 6 |
| *Helicobacter pylori* 26695 | NC_018939.1_1352 | C12 | 1414567 | 1414578 | HpyAXVI-mut1 | 99.8 | RM2 | m6A | CRTTAA | 6 |
| *Helicobacter pylori* 26695 | NC_018939.1_1364 | | | | HpyAII | 100.0 | R2 | NA | GAAGA | NA |
| *Helicobacter pylori* 26695 | NC_018939.1_1365 | | | | M1.HpyAII | 99.6 | M2 | m6A | GAAGA | 5 |
| *Helicobacter pylori* 26695 | NC_018939.1_1366 | | | | M2.HpyAII | 99.7 | M2 | Nm4C | GAAGA | -2 |
| *Helicobacter pylori* 26695 | NC_018939.1_1367 | G10 | 1432276 | 1432285 | M.HpyAXVII | 100.0 | M3 | m6A | TCAG | 3 |
| *Helicobacter pylori* 26695 | NC_018939.1_1368 | | | | M.HpyAXVII | 100.0 | M3 | m6A | TCAG | 3 |
| *Helicobacter pylori* 26695 | NC_018939.1_1369 | | | | Vsp69I | 24.2 | R3 | NA | NA | NA |
| *Helicobacter pylori* 26695 | NC_018939.1_1382 | | | | S.Hpy99XV | 72.8 | S | NA | CTTTANNNNNNCTT, AAGNNNNNNCTT | NA |
| *Helicobacter pylori* 26695 | NC_018939.1_1403 | | | | KpnBI | 47.4 | R1 | NA | CAAANNNNNNRTCA | NA |
| *Helicobacter pylori* 26695 | NC_018939.1_1404 | | | | M.HpyUM032XII | 93.7 | M1 | m6A | CRTANNNNNNNTC | 4 |
| *Helicobacter pylori* 26695 | NC_018939.1_1405 | | | | S.Lla7I | 56.0 | S | NA | NA | NA |
| *Helicobacter pylori* 26695 | NC_018939.1_1471 | G14 | 1544069 | 1544082 | S.Hpy99XXII-mut1 | 69.8 | S | NA | CYANNNNNNTGA | NA |
| *Helicobacter pylori* 26695 | NC_018939.1_1472 | | | | Hpy99XXII | 93.5 | RM2 | m6A | CYANNNNNNTGA | 3 |
| *Helicobacter pylori* 26695 | NC_018939.1_1519 | | | | HpyAXIV | 100.0 | RM2 | m6A | GCGTA | 5 |
| *Helicobacter pylori* 26695 | NC_018939.1_1520 | | | | Hpy99XIII | 82.5 | RM2 | m6A | GCCTA | 5 |

| | | | | | | | | | | |
|---|---|---|---|---|---|---|---|---|---|---|
| *Helicobacter pylori* 26695 | NC_018939.1_1521 | | | | Btr192II | 29.0 | R3 | NA | ACATC | NA |
| *Helicobacter pylori* 26695 | NC_018939.1_1522 | G12 (24 bp upstream) | 1600725 | 1600736 | M.Hpy300X | 93.3 | M3 | m6A | GACY | 2 |
| *Helicobacter pylori* 26695 | NC_018939.1_1523 | | | | M.Hpy99XXI | 65.7 | M3 | m6A | GWCAY | 4 |
| *Helicobacter pylori* 26695 | NC_018939.1_1538 | | | | M.HpyUM032VIII | 100.0 | M2 | m6A | TGCA | 4 |
| *Helicobacter pylori* 3401T | HPY3401T_00730 | | | | S.Hpy66XV | 89.5 | S | NA | CAGNNNNNTC | NA |
| *Helicobacter pylori* 3401T | HPY3401T_00740 | | | | M.Hpy66XV | 94.3 | M1 | m6A | CAGNNNNNTC | 2 |
| *Helicobacter pylori* 3401T | HPY3401T_00750 | | | | CjeFIV | 25.0 | R1 | NA | GCANNNNNRTTA | NA |
| *Helicobacter pylori* 3401T | HPY3401T_00940 | | | | M.HpyUM032VII | 96.9 | M2 | m6A | ATTAAT | 5 |
| *Helicobacter pylori* 3401T | HPY3401T_00970 | | | | M.HpyUM032II | 98.3 | M2 | Nm4C | ACNGT | 2 |
| *Helicobacter pylori* 3401T | HPY3401T_01200 | | | | M.Hpy66XII | 97.3 | M2 | m6A | GTAC | 3 |
| *Helicobacter pylori* 3401T | HPY3401T_02460 | T19 (46 bp upstream) | 260234 | 260252 | EcoCTGmrSD | 23.2 | R4 | NA | NA | NA |
| *Helicobacter pylori* 3401T | HPY3401T_02930 | | | | HpyAXVIIIC | 77.4 | RM2 | NA | GGANNAG | NA |
| *Helicobacter pylori* 3401T | HPY3401T_02940 | | | | HpyAXVIIIC | 74.0 | RM2 | NA | GGANNAG | NA |
| *Helicobacter pylori* 3401T | HPY3401T_02950 | | | | HpyAXVIIIC | 85.6 | RM2 | NA | GGANNAG | NA |
| *Helicobacter pylori* 3401T | HPY3401T_02960 | | | | HpyAXVIIIB | 88.3 | RM2 | NA | GGANNAG | NA |
| *Helicobacter pylori* 3401T | HPY3401T_02970 | | | | HpyAXVIIIB | 80.8 | RM2 | NA | GGANNAG | NA |
| *Helicobacter pylori* 3401T | HPY3401T_02980 | | | | HpyAXVIIIA | 75.7 | RM2 | NA | GGANNAG | NA |
| *Helicobacter pylori* 3401T | HPY3401T_04110 | | | | S.HpyAXIII | 82.1 | S | NA | CTANNNNNNNNTGT | NA |
| *Helicobacter pylori* 3401T | HPY3401T_04660 | A9 | 482749 | 482757 | Hpy99XVI | 91.0 | R1 | NA | RTAYNNNNNRTAY | NA |
| *Helicobacter pylori* 3401T | HPY3401T_04670 | | | | S.HpyAXIII | 85.0 | S | NA | CTANNNNNNNNTGT | NA |
| *Helicobacter pylori* 3401T | HPY3401T_04680 | | | | M.HpyAXIII | 91.7 | M1 | m6A | CTANNNNNNNNTGT | 3 |
| *Helicobacter pylori* 3401T | HPY3401T_04690 | | | | M.NgoAII | 65.9 | M2 | m5C | GGCC | 3 |
| *Helicobacter pylori* 3401T | HPY3401T_04700 | | | | HaeIII | 40.8 | R2 | NA | GGCC | NA |
| *Helicobacter pylori* 3401T | HPY3401T_05220 | | | | M.Hpy66XIII | 97.6 | M2 | m6A | GTNNAC | 5 |
| *Helicobacter pylori* 3401T | HPY3401T_06640 | | | | M.Hpy66VI | 98.4 | M2 | m6A | TCNNGA | 6 |
| *Helicobacter pylori* 3401T | HPY3401T_06650 | | | | HpyGI | 95.1 | R2 | NA | TCNNGA | NA |
| *Helicobacter pylori* 3401T | HPY3401T_06970 | | | | M.HpyUM032XV | 97.8 | M2 | m5C | GCGC | 2 |
| *Helicobacter pylori* 3401T | HPY3401T_07840 | | | | M.HpyUM032I | 97.3 | M2 | m6A | CATG | 2 |
| *Helicobacter pylori* 3401T | HPY3401T_09140 | | | | M1.Hpy66XIV | 94.4 | M2 | m6A | CCTC | -2 |
| *Helicobacter pylori* 3401T | HPY3401T_09240 | | | | HpyAIV | 90.6 | R2 | NA | GANTC | NA |
| *Helicobacter pylori* 3401T | HPY3401T_09250 | | | | HpyAIV | 91.9 | R2 | NA | GANTC | NA |
| *Helicobacter pylori* 3401T | HPY3401T_09260 | | | | M.HpyUM032IV | 99.4 | M2 | m6A | GANTC | 2 |
| *Helicobacter pylori* 3401T | HPY3401T_09270 | C19 | 961250 | 961268 | HpyAXVI-mut2 | 89.8 | RM2 | m6A | CRTCNA | 6 |
| *Helicobacter pylori* 3401T | HPY3401T_09280 | C18 | 962083 | 962100 | HpyAXVI-mut1 | 89.0 | RM2 | m6A | CRTTAA | 6 |
| *Helicobacter pylori* 3401T | HPY3401T_09400 | | | | HpyAII | 90.4 | R2 | NA | GAAGA | NA |
| *Helicobacter pylori* 3401T | HPY3401T_10440 | | | | KpnBI | 43.8 | R1 | NA | CAAANNNNNNRTCA | NA |
| *Helicobacter pylori* 3401T | HPY3401T_10450 | A9 | 1081616 | 1081624 | KpnBI | 50.2 | R1 | NA | CAAANNNNNNRTCA | NA |
| *Helicobacter pylori* 3401T | HPY3401T_10460 | | | | M.HpyUM032XII | 96.4 | M1 | m6A | CRTANNNNNNNTC | 4 |
| *Helicobacter pylori* 3401T | HPY3401T_10470 | | | | M.HpyUM032XII | 94.3 | M1 | m6A | CRTANNNNNNNTC | 4 |
| *Helicobacter pylori* 3401T | HPY3401T_10490 | | | | S.HpyUM032XII | 97.6 | S | NA | CRTANNNNNNNTC | NA |
| *Helicobacter pylori* 3401T | HPY3401T_10600 | | | | M.Hpy99XXI | 76.7 | M3 | m6A | GWCAY | 4 |
| *Helicobacter pylori* 3401T | HPY3401T_10630 | | | | M.Hpy300X | 64.7 | M3 | m6A | GACY | 2 |
| *Helicobacter pylori* 3401T | HPY3401T_10640 | | | | Btr192II | 29.8 | R3 | NA | ACATC | NA |
| *Helicobacter pylori* 3401T | HPY3401T_10660 | | | | Hpy99XIII | 80.2 | RM2 | m6A | GCCTA | 5 |
| *Helicobacter pylori* 3401T | HPY3401T_10670 | | | | Hpy99XIII | 83.1 | RM2 | m6A | GCCTA | 5 |
| *Helicobacter pylori* 3401T | HPY3401T_10830 | | | | HpyC1I | 95.8 | R2 | NA | CCATC | NA |
| *Helicobacter pylori* 3401T | HPY3401T_10840 | | | | M2.Hpy66X | 99.3 | M2 | m6A | CCATC | -2 |
| *Helicobacter pylori* 3401T | HPY3401T_10870 | | | | M2.Hpy300VI | 98.9 | M2 | m6A | CCATC | 3 |
| *Helicobacter pylori* 3401T | HPY3401T_10880 | | | | M1.Hpy66X | 98.5 | M2 | m6A | CCATC | 3 |
| *Helicobacter pylori* 3401T | HPY3401T_11150 | | | | HpyUM032XIII | 97.2 | RM2 | m6A | CYANNNNNNNTRG | 3 |
| *Helicobacter pylori* 3401T | HPY3401T_11160 | | | | S.Hpy99XXII-mut1 | 43.4 | S | NA | CYANNNNNNTGA | NA |
| *Helicobacter pylori* 3401T | HPY3401T_11810 | C17 | 1230764 | 1230780 | LlaFI | 29.6 | R3 | NA | NA | NA |
| *Helicobacter pylori* 3401T | HPY3401T_11820 | | | | M.LlaFI | 39.1 | M3 | m6A | ? | ? |
| *Helicobacter pylori* 3401T | HPY3401T_12070 | | | | M1.Hpy66XIV | 96.2 | M2 | m6A | CCTC | -2 |
| *Helicobacter pylori* 3401T | HPY3401T_12080 | | | | M1.HpyUM032VI | 98.9 | M2 | m6A | CCTC | -2 |
| *Helicobacter pylori* 3401T | HPY3401T_12090 | | | | M.HpyD27I | 97.9 | M2 | m5C | CCTC | 1 |
| *Helicobacter pylori* 3401T | HPY3401T_12920 | | | | M.HpyUM032III | 97.1 | M2 | m6A | GATC | 2 |
| *Helicobacter pylori* 3401T | HPY3401T_12930 | | | | HpyAIII | 89.9 | R2 | NA | GATC | NA |
| *Helicobacter pylori* 3401T | HPY3401T_13500 | | | | R1.LlaJI | 47.1 | R2 | NA | NA | NA |

| | | | | | | | | | | |
|---|---|---|---|---|---|---|---|---|---|---|
| *Helicobacter pylori* 3401T | HPY3401T_13510 | | | | R2.LlaJI | 41.1 | R2 | NA | NA | NA |
| *Helicobacter pylori* 3401T | HPY3401T_14360 | | | | M.HpyUM032XVII | 98.2 | M2 | m6A | TCGA | 4 |
| *Helicobacter pylori* 3401T | HPY3401T_14400 | | | | M.Hpy99VIII | 98.9 | M2 | Nm4C | CCGG | 1 |
| *Helicobacter pylori* 3401T | HPY3401T_14410 | | | | M.HpyUM032IX | 98.1 | M2 | Nm4C | CCGG | 1 |
| *Helicobacter pylori* 3401T | HPY3401T_14890 | | | | HpyHI | 93.9 | R2 | NA | CTNAG | NA |
| *Helicobacter pylori* HPK5 | HPYHPK5_00020 | | | | KpnBI | 48.1 | R1 | NA | CAAANNNNNNRTCA | NA |
| *Helicobacter pylori* HPK5 | HPYHPK5_00030 | | | | M.HpyUM032XII | 96.9 | M1 | m6A | CRTANNNNNNNTC | 4 |
| *Helicobacter pylori* HPK5 | HPYHPK5_00040 | | | | M.HpyUM032XII | 93.5 | M1 | m6A | CRTANNNNNNNTC | 4 |
| *Helicobacter pylori* HPK5 | HPYHPK5_00060 | | | | S.HpyUM032XII | 70.9 | S | NA | CRTANNNNNNNTC | NA |
| *Helicobacter pylori* HPK5 | HPYHPK5_00170 | G9 | 20576 | 20584 | M.Hpy300X | 69.3 | M3 | m6A | GACY | 2 |
| *Helicobacter pylori* HPK5 | HPYHPK5_00180 | | | | M.Hpy99XXI | 91.2 | M3 | m6A | GWCAY | 4 |
| *Helicobacter pylori* HPK5 | HPYHPK5_00190 | | | | Btr192II | 30.0 | R3 | NA | ACATC | NA |
| *Helicobacter pylori* HPK5 | HPYHPK5_00200 | | | | HpyAXIV | 74.7 | RM2 | m6A | GCGTA | 5 |
| *Helicobacter pylori* HPK5 | HPYHPK5_00360 | | | | HpyC1I | 96.3 | R2 | NA | CCATC | NA |
| *Helicobacter pylori* HPK5 | HPYHPK5_00370 | | | | M2.Hpy66X | 98.7 | M2 | m6A | CCATC | -2 |
| *Helicobacter pylori* HPK5 | HPYHPK5_00380 | | | | M1.Hpy66X | 99.4 | M2 | m6A | CCATC | 3 |
| *Helicobacter pylori* HPK5 | HPYHPK5_00650 | | | | BtsCI | 33.8 | R2 | NA | GGATG | NA |
| *Helicobacter pylori* HPK5 | HPYHPK5_00660 | | | | M4.BstF5I | 54.7 | M2 | m6A | GGATG | 3 |
| *Helicobacter pylori* HPK5 | HPYHPK5_00670 | | | | M.BtsCI | 51.0 | M2 | NA | GGATG | NA |
| *Helicobacter pylori* HPK5 | HPYHPK5_01220 | | | | EcoRI | 67.4 | R2 | NA | GAATTC | NA |
| *Helicobacter pylori* HPK5 | HPYHPK5_01230 | | | | EcoRI | 55.6 | R2 | NA | GAATTC | NA |
| *Helicobacter pylori* HPK5 | HPYHPK5_01240 | | | | M.Hpy300III | 95.9 | M2 | m6A | GAATTC | 3 |
| *Helicobacter pylori* HPK5 | HPYHPK5_01570 | | | | LlaFI | 29.1 | R3 | NA | NA | NA |
| *Helicobacter pylori* HPK5 | HPYHPK5_01580 | | | | M.LlaFI | 38.9 | M3 | m6A | ? | ? |
| *Helicobacter pylori* HPK5 | HPYHPK5_03110 | | | | M.HpyUM032I | 97.3 | M2 | m6A | CATG | 2 |
| *Helicobacter pylori* HPK5 | HPYHPK5_03970 | | | | M.HpyUM032XV | 94.2 | M2 | m5C | GCGC | 2 |
| *Helicobacter pylori* HPK5 | HPYHPK5_04300 | | | | HpyGI | 96.5 | R2 | NA | TCNNGA | NA |
| *Helicobacter pylori* HPK5 | HPYHPK5_04310 | | | | M.Hpy66VI | 95.4 | M2 | m6A | TCNNGA | 6 |
| *Helicobacter pylori* HPK5 | HPYHPK5_05630 | | | | M.HpyUM032XI | 97.1 | M2 | m6A | GTNNAC | 5 |
| *Helicobacter pylori* HPK5 | HPYHPK5_06160 | | | | M.HpyAXIII | 90.5 | M1 | m6A | CTANNNNNNNNTGT | 3 |
| *Helicobacter pylori* HPK5 | HPYHPK5_06170 | | | | S.HpyAXIII | 94.8 | S | NA | CTANNNNNNNNTGT | NA |
| *Helicobacter pylori* HPK5 | HPYHPK5_06180 | | | | Hpy99XVI | 91.4 | R1 | NA | RTAYNNNNNRTAY | NA |
| *Helicobacter pylori* HPK5 | HPYHPK5_06730 | | | | S.HpyAXIII | 94.8 | S | NA | CTANNNNNNNNTGT | NA |
| *Helicobacter pylori* HPK5 | HPYHPK5_07870 | | | | HpyAXVIIIC | 83.3 | RM2 | NA | GGANNAG | NA |
| *Helicobacter pylori* HPK5 | HPYHPK5_07880 | | | | HpyAXVIIIA | 75.7 | RM2 | NA | GGANNAG | NA |
| *Helicobacter pylori* HPK5 | HPYHPK5_09400 | | | | EcoCTGmrSD | 25.3 | R4 | NA | NA | NA |
| *Helicobacter pylori* HPK5 | HPYHPK5_09680 | | | | M.HpyUM032II | 97.8 | M2 | Nm4C | ACNGT | 2 |
| *Helicobacter pylori* HPK5 | HPYHPK5_09710 | | | | M.HpyUM032VII | 97.6 | M2 | m6A | ATTAAT | 5 |
| *Helicobacter pylori* HPK5 | HPYHPK5_09880 | C13 | 1055137 | 1055149 | PhaAI | 31.0 | R1 | NA | NA | NA |
| *Helicobacter pylori* HPK5 | HPYHPK5_09890 | | | | CjeFIV | 24.8 | R1 | NA | GCANNNNNRTTA | NA |
| *Helicobacter pylori* HPK5 | HPYHPK5_09900 | | | | M.Hpy66XV | 95.0 | M1 | m6A | CAGNNNNNTC | 2 |
| *Helicobacter pylori* HPK5 | HPYHPK5_09910 | | | | S.Hpy99XX | 85.5 | S | NA | CCANNNNNNTTT | NA |
| *Helicobacter pylori* HPK5 | HPYHPK5_11150 | | | | M.TmaI | 38.2 | M2 | Nm4C | CGCG | ? |
| *Helicobacter pylori* HPK5 | HPYHPK5_11160 | | | | HpyHI | 92.6 | R2 | NA | CTNAG | NA |
| *Helicobacter pylori* HPK5 | HPYHPK5_11660 | | | | M.Hpy99VIII | 98.8 | M2 | Nm4C | CCGG | 1 |
| *Helicobacter pylori* HPK5 | HPYHPK5_12500 | | | | R2.LlaJI | 41.0 | R2 | NA | NA | NA |
| *Helicobacter pylori* HPK5 | HPYHPK5_12510 | | | | R1.LlaJI | 51.3 | R2 | NA | NA | NA |
| *Helicobacter pylori* HPK5 | HPYHPK5_12530 | | | | M.Cce743I | 41.4 | M2 | m5C | GACGC | 5 |
| *Helicobacter pylori* HPK5 | HPYHPK5_12540 | | | | M.Cce743I | 44.8 | M2 | m5C | GACGC | 5 |
| *Helicobacter pylori* HPK5 | HPYHPK5_13360 | | | | M.Hpy66III | 98.2 | M2 | m6A | GATC | 2 |
| *Helicobacter pylori* HPK5 | HPYHPK5_13370 | | | | HpyAIII | 88.4 | R2 | NA | GATC | NA |
| *Helicobacter pylori* HPK5 | HPYHPK5_13790 | | | | M.HpyAV | 84.6 | M2 | m5C | CCTTC | 2 |
| *Helicobacter pylori* HPK5 | HPYHPK5_13810 | | | | M.HpyD27I | 96.6 | M2 | m5C | CCTC | 1 |
| *Helicobacter pylori* HPK5 | HPYHPK5_13820 | | | | M1.Hpy66XIV | 96.5 | M2 | m6A | CCTC | -2 |
| *Helicobacter pylori* HPK5 | HPYHPK5_13920 | C14 | 1456590 | 1456603 | Hpy300XI | 88.2 | RM2 | m6A | CCTYNA | 6 |
| *Helicobacter pylori* HPK5 | HPYHPK5_13930 | C15 | 1457409 | 1457423 | HpyAXVI-mut1 | 87.3 | RM2 | m6A | CRTTAA | 6 |
| *Helicobacter pylori* HPK5 | HPYHPK5_14050 | | | | HpyAII | 92.4 | R2 | NA | GAAGA | NA |
| *Helicobacter pylori* HPK5 | HPYHPK5_14060 | | | | HpyAII | 96.8 | R2 | NA | GAAGA | NA |
| *Helicobacter pylori* HPK5 | HPYHPK5_14070 | | | | M1.HpyAII | 96.9 | M2 | m6A | GAAGA | 5 |
| *Helicobacter pylori* HPK5 | HPYHPK5_14080 | | | | M2.HpyAII | 96.9 | M2 | Nm4C | GAAGA | -2 |

| | | | | | | | |
|---|---|---|---|---|---|---|---|
| *Helicobacter pylori* HPK5 | HPYHPK5_14090 | M.HpyAXVII | 71.3 | M3 | m6A | TCAG | 3 |
| *Helicobacter pylori* HPK5 | HPYHPK5_14100 | Vsp69I | 25.0 | R3 | NA | NA | NA |
| *Helicobacter pylori* HPK5 | HPYHPK5_14970 | M.HpyUM032VIII | 98.7 | M2 | m6A | TGCA | 4 |

**Table S3. Predicted phage defense systems in *H. pylori* strains.**

| System number | Strain | Predicted defence system | CDS ID |
|---|---|---|---|
| Abi-1 | *Helicobacter pylori* 26695 | AbiL | NC_018939.1_1077 |
| Abi-1 | *Helicobacter pylori* 26695 | AbiL | NC_018939.1_1076 |
| RM-1 | *Helicobacter pylori* 26695 | RM_type_I | NC_018939.1_459 |
| RM-1 | *Helicobacter pylori* 26695 | RM_type_I | NC_018939.1_458 |
| RM-1 | *Helicobacter pylori* 26695 | RM_type_I | NC_018939.1_457 |
| RM-2 | *Helicobacter pylori* 26695 | RM_type_I | NC_018939.1_845 |
| RM-2 | *Helicobacter pylori* 26695 | RM_type_I | NC_018939.1_843 |
| RM-2 | *Helicobacter pylori* 26695 | RM_type_I | NC_018939.1_841 |
| RM-3 | *Helicobacter pylori* 26695 | RM_type_I | NC_018939.1_1395 |
| RM-3 | *Helicobacter pylori* 26695 | RM_type_I | NC_018939.1_1394 |
| RM-4 | *Helicobacter pylori* 26695 | RM_type_II | NC_018939.1_49 |
| RM-4 | *Helicobacter pylori* 26695 | RM_type_II | NC_018939.1_48 |
| RM-4 | *Helicobacter pylori* 26695 | RM_type_II | NC_018939.1_47 |
| RM-4 | *Helicobacter pylori* 26695 | RM_type_II | NC_018939.1_45 |
| RM-4 | *Helicobacter pylori* 26695 | RM_type_II | NC_018939.1_44 |
| RM-5 | *Helicobacter pylori* 26695 | RM_type_II | NC_018939.1_87 |
| RM-5 | *Helicobacter pylori* 26695 | RM_type_II | NC_018939.1_86 |
| RM-6 | *Helicobacter pylori* 26695 | RM_type_II | NC_018939.1_361 |
| RM-6 | *Helicobacter pylori* 26695 | RM_type_II | NC_018939.1_360 |
| RM-7 | *Helicobacter pylori* 26695 | RM_type_II | NC_018939.1_503 |
| RM-7 | *Helicobacter pylori* 26695 | RM_type_II | NC_018939.1_501 |
| RM-8 | *Helicobacter pylori* 26695 | RM_type_II | NC_018939.1_1344 |
| RM-8 | *Helicobacter pylori* 26695 | RM_type_II | NC_018939.1_1343 |
| RM-9 | *Helicobacter pylori* 26695 | RM_type_II | NC_018939.1_1360 |
| RM-9 | *Helicobacter pylori* 26695 | RM_type_II | NC_018939.1_1359 |
| RM-9 | *Helicobacter pylori* 26695 | RM_type_II | NC_018939.1_1358 |
| RM-10 | *Helicobacter pylori* 26695 | RM_type_IIG | NC_018939.1_670 |
| RM-11 | *Helicobacter pylori* 26695 | RM_type_IIG | NC_018939.1_1461 |
| RM-12 | *Helicobacter pylori* 26695 | RM_type_IIG | NC_018939.1_1505 |
| RM-13 | *Helicobacter pylori* 26695 | RM_type_III | NC_018939.1_593 |
| RM-13 | *Helicobacter pylori* 26695 | RM_type_III | NC_018939.1_592 |
| RM-14 | *Helicobacter pylori* 26695 | RM_type_III | NC_018939.1_1511 |
| RM-14 | *Helicobacter pylori* 26695 | RM_type_III | NC_018939.1_1509 |
| Abi-1 | *Helicobacter pylori* 3401T | AbiL | HPY3401T_00050 |
| Abi-1 | *Helicobacter pylori* 3401T | AbiL | HPY3401T_00040 |
| RM-1 | *Helicobacter pylori* 3401T | RM_type_I | HPY3401T_04680 |
| RM-1 | *Helicobacter pylori* 3401T | RM_type_I | HPY3401T_04670 |
| RM-1 | *Helicobacter pylori* 3401T | RM_type_I | HPY3401T_04660 |
| RM-2 | *Helicobacter pylori* 3401T | RM_type_I | HPY3401T_10490 |
| RM-2 | *Helicobacter pylori* 3401T | RM_type_I | HPY3401T_10460 |
| RM-2 | *Helicobacter pylori* 3401T | RM_type_I | HPY3401T_10450 |
| RM-2 | *Helicobacter pylori* 3401T | RM_type_II | HPY3401T_04700 |
| RM-2 | *Helicobacter pylori* 3401T | RM_type_II | HPY3401T_04690 |
| RM-2 | *Helicobacter pylori* 3401T | RM_type_II | HPY3401T_04680 |
| RM-3 | *Helicobacter pylori* 3401T | RM_type_II | HPY3401T_06650 |
| RM-3 | *Helicobacter pylori* 3401T | RM_type_II | HPY3401T_06640 |
| RM-4 | *Helicobacter pylori* 3401T | RM_type_II | HPY3401T_09260 |
| RM-4 | *Helicobacter pylori* 3401T | RM_type_II | HPY3401T_09240 |
| RM-5 | *Helicobacter pylori* 3401T | RM_type_II | HPY3401T_10840 |
| RM-5 | *Helicobacter pylori* 3401T | RM_type_II | HPY3401T_10830 |
| RM-6 | *Helicobacter pylori* 3401T | RM_type_II | HPY3401T_12930 |
| RM-6 | *Helicobacter pylori* 3401T | RM_type_II | HPY3401T_12920 |
| RM-7 | *Helicobacter pylori* 3401T | RM_type_IIG | HPY3401T_11150 |
| RM-8 | *Helicobacter pylori* 3401T | RM_type_III | HPY3401T_10640 |
| RM-8 | *Helicobacter pylori* 3401T | RM_type_III | HPY3401T_10630 |
| RM-9 | *Helicobacter pylori* 3401T | RM_type_III | HPY3401T_11820 |
| RM-9 | *Helicobacter pylori* 3401T | RM_type_III | HPY3401T_11810 |
| RM-1 | *Helicobacter pylori* HPK5 | RM_type_I | HPYHPK5_06160 |
| RM-1 | *Helicobacter pylori* HPK5 | RM_type_I | HPYHPK5_06150 |
| RM-1 | *Helicobacter pylori* HPK5 | RM_type_I | HPYHPK5_06140 |
| RM-2 | *Helicobacter pylori* HPK5 | RM_type_II | HPYHPK5_00360 |
| RM-2 | *Helicobacter pylori* HPK5 | RM_type_II | HPYHPK5_00350 |
| RM-2 | *Helicobacter pylori* HPK5 | RM_type_II | HPYHPK5_00340 |
| RM-3 | *Helicobacter pylori* HPK5 | RM_type_II | HPYHPK5_00650 |
| RM-3 | *Helicobacter pylori* HPK5 | RM_type_II | HPYHPK5_00640 |
| RM-3 | *Helicobacter pylori* HPK5 | RM_type_II | HPYHPK5_00630 |
| RM-4 | *Helicobacter pylori* HPK5 | RM_type_II | HPYHPK5_01220 |
| RM-4 | *Helicobacter pylori* HPK5 | RM_type_II | HPYHPK5_01200 |
| RM-5 | *Helicobacter pylori* HPK5 | RM_type_II | HPYHPK5_04290 |
| RM-5 | *Helicobacter pylori* HPK5 | RM_type_II | HPYHPK5_04280 |
| RM-6 | *Helicobacter pylori* HPK5 | RM_type_II | HPYHPK5_11140 |
| RM-6 | *Helicobacter pylori* HPK5 | RM_type_II | HPYHPK5_11130 |
| RM-7 | *Helicobacter pylori* HPK5 | RM_type_II | HPYHPK5_12510 |
| RM-7 | *Helicobacter pylori* HPK5 | RM_type_II | HPYHPK5_12490 |
| RM-7 | *Helicobacter pylori* HPK5 | RM_type_II | HPYHPK5_12480 |
| RM-8 | *Helicobacter pylori* HPK5 | RM_type_II | HPYHPK5_13350 |
| RM-8 | *Helicobacter pylori* HPK5 | RM_type_II | HPYHPK5_13340 |
| RM-9 | *Helicobacter pylori* HPK5 | RM_type_II | HPYHPK5_13800 |
| RM-9 | *Helicobacter pylori* HPK5 | RM_type_II | HPYHPK5_13790 |
| RM-9 | *Helicobacter pylori* HPK5 | RM_type_II | HPYHPK5_13780 |
| RM-10 | *Helicobacter pylori* HPK5 | RM_type_II | HPYHPK5_14070 |
| RM-10 | *Helicobacter pylori* HPK5 | RM_type_II | HPYHPK5_14060 |
| RM-10 | *Helicobacter pylori* HPK5 | RM_type_II | HPYHPK5_14050 |
| RM-10 | *Helicobacter pylori* HPK5 | RM_type_II | HPYHPK5_14040 |
| RM-11 | *Helicobacter pylori* HPK5 | RM_type_IIG | HPYHPK5_00180 |
| RM-12 | *Helicobacter pylori* HPK5 | RM_type_IIG | HPYHPK5_07850 |
| RM-13 | *Helicobacter pylori* HPK5 | RM_type_III | HPYHPK5_00170 |
| RM-13 | *Helicobacter pylori* HPK5 | RM_type_III | HPYHPK5_00150 |
| RM-14 | *Helicobacter pylori* HPK5 | RM_type_III | HPYHPK5_01560 |
| RM-14 | *Helicobacter pylori* HPK5 | RM_type_III | HPYHPK5_01550 |

**Table S4. Methylated motifs predicted from *H. pylori* strains.**

| Strain | Motif sequence | Center possition | Modification type | Number of methylated sites | Number of motif sequences | Methylation ratio (%) | Reference |
|---|---|---|---|---|---|---|---|
| *Helicobacter pylori* 26695 | ACANNNNNNNNTAG | 3 | m6A | 326 | 330 | 98.8 | Krebes et al. Nucleic Acids Res. (2014) |
| *Helicobacter pylori* 26695 | CTANNNNNNNNTGT | 3 | m6A | 326 | 330 | 98.8 | Krebes et al. Nucleic Acids Res. (2014) |
| *Helicobacter pylori* 26695 | ATTAAT | 5 | m6A | 957 | 972 | 98.5 | Krebes et al. Nucleic Acids Res. (2014) |
| *Helicobacter pylori* 26695 | CATG | 2 | m6A | 14741 | 14782 | 99.7 | Krebes et al. Nucleic Acids Res. (2014) |
| *Helicobacter pylori* 26695 | CCTC | 1 | m5C | 3913 | 4885 | 80.1 | Krebes et al. Nucleic Acids Res. (2014) |
| *Helicobacter pylori* 26695 | GAGG | 2 | m6A | 4841 | 4885 | 99.1 | Krebes et al. Nucleic Acids Res. (2014) |
| *Helicobacter pylori* 26695 | CCTTC | 2 | m5C | 1198 | 1304 | 91.9 | Krebes et al. Nucleic Acids Res. (2014) |
| *Helicobacter pylori* 26695 | GAAGG | 3 | m6A | 1252 | 1304 | 96.0 | Krebes et al. Nucleic Acids Res. (2014) |
| *Helicobacter pylori* 26695 | CGRAT | 4 | m6A | 3240 | 3275 | 98.9 | Krebes et al. Nucleic Acids Res. (2014) |
| *Helicobacter pylori* 26695 | GAAGA | 5 | m6A | 4809 | 4823 | 99.7 | Krebes et al. Nucleic Acids Res. (2014) |
| *Helicobacter pylori* 26695 | TCTTC | 2 | m4C | 4593 | 4823 | 95.2 | Krebes et al. Nucleic Acids Res. (2014) |
| *Helicobacter pylori* 26695 | GANTC | 2 | m6A | 5549 | 5570 | 99.6 | Krebes et al. Nucleic Acids Res. (2014) |
| *Helicobacter pylori* 26695 | GATC | 2 | m6A | 10764 | 10834 | 99.4 | Krebes et al. Nucleic Acids Res. (2014) |
| *Helicobacter pylori* 26695 | GCAG | 3 | m6A | 4234 | 4260 | 99.4 | Krebes et al. Nucleic Acids Res. (2014) |
| *Helicobacter pylori* 26695 | GCGC | 2 | m5C | 6328 | 12538 | 50.5 | Krebes et al. Nucleic Acids Res. (2014) |
| *Helicobacter pylori* 26695 | GCGTA | 5 | m6A | 2046 | 2058 | 99.4 | Krebes et al. Nucleic Acids Res. (2014) |
| *Helicobacter pylori* 26695 | TCGA | 4 | m6A | 601 | 608 | 98.8 | Krebes et al. Nucleic Acids Res. (2014) |
| *Helicobacter pylori* 3401T | CATG | 2 | m6A | 13976 | 14288 | 97.8 | This study |
| *Helicobacter pylori* 3401T | GATC | 2 | m6A | 9821 | 9976 | 98.4 | This study |
| *Helicobacter pylori* 3401T | GANTC | 2 | m6A | 5371 | 5458 | 98.4 | This study |
| *Helicobacter pylori* 3401T | CGRA | 4 | m6A | 7018 | 8262 | 84.9 | This study |
| *Helicobacter pylori* 3401T | TCNNGA | 6 | m6A | 3715 | 3826 | 97.1 | This study |
| *Helicobacter pylori* 3401T | CTANNNNNNNNTTG | 3 | m6A | 815 | 829 | 98.3 | This study |
| *Helicobacter pylori* 3401T | CAANNNNNNNNTAG | 3 | m6A | 817 | 829 | 98.6 | This study |
| *Helicobacter pylori* 3401T | ATTAAT | 5 | m6A | 814 | 874 | 93.1 | This study |
| *Helicobacter pylori* 3401T | GTNNAC | 5 | m6A | 490 | 502 | 97.6 | This study |
| *Helicobacter pylori* 3401T | GATGG | 2 | m6A | 1623 | 2234 | 72.6 | This study |
| *Helicobacter pylori* 3401T | CCGCAT | 5 | m6A | 631 | 678 | 93.1 | This study |
| *Helicobacter pylori* 3401T | TCGA | 4 | m6A | 492 | 494 | 99.6 | This study |
| *Helicobacter pylori* 3401T | GTAC | 3 | m6A | 185 | 186 | 99.5 | This study |
| *Helicobacter pylori* 3401T | GACTC | 3 | m5C | 288 | 837 | 34.4 | This study |
| *Helicobacter pylori* 3401T | GCGCGC | 2 | m5C | 259 | 518 | 50.0 | This study |
| *Helicobacter pylori* HPK5 | CATG | 2 | m6A | 13883 | 14172 | 98.0 | This study |
| *Helicobacter pylori* HPK5 | GATC | 2 | m6A | 9883 | 10034 | 98.5 | This study |
| *Helicobacter pylori* HPK5 | TGCA | 4 | m6A | 11143 | 11368 | 98.0 | This study |
| *Helicobacter pylori* HPK5 | CTNAG | 1 | m4C | 4855 | 5530 | 87.8 | This study |
| *Helicobacter pylori* HPK5 | GAKGB | 2 | m6A | 7172 | 10199 | 70.3 | This study |
| *Helicobacter pylori* HPK5 | GAAGA | 5 | m6A | 4253 | 4544 | 93.6 | This study |
| *Helicobacter pylori* HPK5 | TCTTC | 2 | m4C | 4017 | 4544 | 88.4 | This study |
| *Helicobacter pylori* HPK5 | GGCAA | 5 | m6A | 3371 | 3421 | 98.5 | This study |
| *Helicobacter pylori* HPK5 | CCATC | 3 | m6A | 2241 | 2284 | 98.1 | This study |
| *Helicobacter pylori* HPK5 | CCGG | 1 | m4C | 2813 | 3290 | 85.5 | This study |
| *Helicobacter pylori* HPK5 | CATCC | 2 | m6A | 1271 | 1295 | 98.1 | This study |
| *Helicobacter pylori* HPK5 | GAGGA | 2 | m6A | 742 | 764 | 97.1 | This study |
| *Helicobacter pylori* HPK5 | GGATGA | 3 | m6A | 367 | 383 | 95.8 | This study |
| *Helicobacter pylori* HPK5 | CCTTGA | 6 | m6A | 1073 | 1097 | 97.8 | This study |
| *Helicobacter pylori* HPK5 | CTANNNNNNNNTGT | 3 | m6A | 305 | 312 | 97.8 | This study |
| *Helicobacter pylori* HPK5 | GTNNAC | 5 | m6A | 479 | 488 | 98.2 | This study |
| *Helicobacter pylori* HPK5 | GAATTC | 3 | m6A | 294 | 302 | 97.4 | This study |
| *Helicobacter pylori* HPK5 | ACANNNNNNNNTAG | 3 | m6A | 305 | 312 | 97.8 | This study |
| *Helicobacter pylori* HPK5 | CGWAG | 4 | m6A | 1103 | 1804 | 61.1 | This study |
| *Helicobacter pylori* HPK5 | GCGCGC | 2 | m5C | 221 | 520 | 42.5 | This study |

#### Transparent Methods

##### Culture conditions of *H. pylori* and phages

*H. pylori* 26695, 3401T (derived from *H. pylori* 3401), and HPK5 were routinely cultured on BEV plates composed of Brucella broth (Becton Dickinson, Cockeysville, MD, USA) supplemented with 1.5% agar, 10% heat-inactivated equine serum (Hyclone Laboratories, South Logan, UT, USA), and 10 μg/mL vancomycin. Cultures were incubated at 37 °C under microaerobic conditions (10% $CO_2$). For precultures, *H. pylori* was incubated in 5 mL of BEV broth at 37 °C under microaerobic conditions with shaking at 200 rpm reciprocating shaking using an EYEKA multishaker (Tokyo Rikakikai, Tokyo, Japan) for 20 h.

Phage KHP30T stock was prepared from the filtered supernatant of *H. pylori* 3401 after inoculation with KHP30, as described previously.[1] Phage stocks were stored at 4 °C. Nine adapted phage cultures were generated using KHP30T, infecting three *H. pylori* strains (26695, 3401T, and HPK5) in sequential steps (Figure 1). Detailed procedures for phage adaptation are described in the following section.

##### Preparation of adapted phages

In the first stage of the multistage infection procedure (Figure 1), KHP30T was adapted to *H. pylori* 26695. First, 100 μL of KHP30T stock (approximately $1\times10^7$ PFU/mL) and 500 μL of *H. pylori* 26695 preculture (approximately $1\times10^8$ colony forming units [CFU]/mL) were co-inoculated on BEV plates layered with top agar (7 mL of Brucella broth supplemented with 0.5% agar). After incubation of the double-layered plate at 37 °C under microaerobic conditions for 3 days, a single clear lytic plaque was picked, suspended in 200 μL of BEV broth, and used as a single-plaque suspension. Next, 100 μL of the single-plaque suspension and 500 μL of *H. pylori* 26695 preculture were co-cultured on double-layered plates at 37 °C under microaerobic conditions for 3 days. Upon confirming confluent lysis, the top agar was collected in 3–4 mL of BEV broth and centrifuged (10,000 ×*g*, 10 min, 4 °C). The supernatant was passed through a 0.45-μm syringe filter and used as a phage solution. Finally, 100 μL of phage solution and 500 μL of *H. pylori* 26695 preculture were co-cultured in 5 double-layer plates at 37°C under microaerobic conditions for 3 days. The top agar was collected, centrifuged (10,000 ×*g*, 10 min, 4°C) and passed through a 0.45-μm syringe filter. The resulting solution was used as an adapted phage stock and named K2 (1st generation in Figure 1). In the second stage, the adapted phage stock of K2 was co-cultured with *H. pylori* 3401T and HPK5 in the same manner as described above to prepare single-plaque suspensions, phage solutions, and adapted phage stocks (Figure S1) of K23 and K2H (2nd stage in Figure 1). In the third generation, K23 was adapted to *H. pylori* 26695 and HPK5 to produce K232 and K23H, respectively, while K2H was adapted to *H. pylori* 26695 and 3401T to yield K2H2 and K2H3, respectively. Single-plaque suspensions, phage solutions, and adapted phages were prepared as described above. In the fourth generation, K23H and K2H3 were adapted to *H. pylori* 3401T and HPK5 to obtain K23H3 and K2H3H, respectively. All adapted phages were stored at 4 °C.

**Titration of adapted phages**

The infectious titers of nine adapted phages (K2, K23, K232, K23H, K23H3, K2H, K2H2, K2H3, and K2H3H) and three reproliferated phages (K2, K23, and K2H, cultured on a larger scale) were measured using a plaque assay.[2] Briefly, 100 µL of each serially 10-fold diluted adapted phage (Figure S1) and 500 µL of *H. pylori* preculture (approximately $1 \times 10^8$ CFU/mL) were co-cultured on double-layered plates. Plaques were counted after incubation at 37 °C under microaerobic conditions for 3 days. Each experiment was performed with three independent biological replicates ($n$ = 3), with each biological replicate including two technical replicates. To test the differences in phage titers, the Tukey−Kramer test was applied using the 'multcomp' R package (v1.4-20).

**Large-scale culture and purification of phages**

Three adapted phages (K2, K23, and K2H) were selected for epigenomic analysis. To extract high-yield (>1 µg) genomic DNA for SMRT sequencing, large-scale cultures and purifications were conducted according to a previous report with minor modifications.[2] Briefly, adapted phage stocks of K2, K23, and K2H were each co-cultured with a preculture of *H. pylori* 26695, 3401T, and HPK5, respectively, in 1 L of BCV medium (Brucella broth supplemented with 0.5% β-cyclodextrin [Nacalai Tesque, Kyoto, Japan] and 10 µg/mL vancomycin) at a multiplicity of infection of 1–5. After incubation at 37 °C under microaerobic conditions with shaking (180 rpm of reciprocating shaking with the EYEKA multishaker) for 3 days, the cell lysate was centrifuged at 10,000 ×*g* at

4 °C for 5 min to remove cell debris, and the supernatant was subjected to polyethyleneglycol (PEG) precipitation. The resulting supernatant was added to 10% (*w/v*) PEG 6000 (Fujifilm Wako Pure Chemical, Osaka, Japan), 3 M NaCl (Nacalai Tesque), and 1% (*v/v*) Tween 20 (Fujifilm Wako Pure Chemical) and incubated at 4 °C for 3 days. After centrifugation at 10,000 ×*g* for 20 min at 4 °C, the pellet was dissolved in 2 mL of TM buffer (10 mM Tris-HCl [pH 7.5] and 5 mM $MgCl_2$) and treated with 100 μg/mL DNase I (Nippon Gene, Tokyo, Japan) and 100 μg/mL RNase A (Nippon Gene) at 37 °C for 30 min. Phage particles concentrated by PEG precipitation were purified by CsCl density gradient ultracentrifugation. The phage solution (2 mL) was layered on top of a discontinuous CsCl density gradient, comprising 1.5, 1.0, and 0.3 mL CsCl solutions (densities 1.3, 1.5, and 1.7, respectively), in a polycarbonate ultracentrifuge tube (Himac S408829A, Eppendorf Himac Technologies, Ibaraki, Japan). The tubes were centrifuged (100,000 ×*g*, 1 h, 4 °C) using a Himac CS150FNX ultracentrifuge, and the phage band (1 mL) was retrieved. The bands were suspended in AAS buffer (100 mM ammonium acetate, 10 mM NaCl, 1 mM $MgCl_2$, and 1 mM $CaCl_2$) and ultracentrifuged (100,000 ×*g*, 1 h, 4 °C) to pellet the phage particles. The resulting purified phage pellet was stored at -80 °C. Titration of large-scale cultured phages was performed as described above to evaluate any potential changes in infection efficiency across the adapted phage cultures during large-scale cultivation.

**DNA extraction and SMRT sequencing**

Genomic DNA of *H. pylori* 3401T and HPK5 was extracted using QIAGEN Genomic-tips 100/G (QIAGEN, Hilden, Germany), according to the manufacturer’s instructions. For SMRT sequencing, library preparation and HiFi sequencing were performed by Macrogen (Tokyo, Japan) on the PacBio Sequel II system (Pacific Biosciences, Menlo Park, CA, USA).

Genomic DNA from three adapted phages (K2H, K2, and K23) was prepared using the phenol-chloroform extraction method after large-scale cultivation and particle purification, as described above. For K2H, the extracted genomic DNA was sheared to 10 kb for SMRTbell library preparation and subjected to SMRT sequencing in continuous long-read mode on the PacBio Sequel system (Pacific Biosciences). For K2 and K23, DNA was sheared to 10–15 kb for SMRTbell library preparation and sequenced in circular consensus sequencing (CCS) mode on the PacBio Sequel II system (Pacific Biosciences). All library preparation and phage sequencing were performed by Azenta (South Plainfield, NJ, USA).

**Bioinformatics**

For *H. pylori* 3401T and HPK5, the HiFi reads were processed to include conventional kinetic information using the ccs-kinetics-by-strandify script provided by PacBio. The reads were assembled *de novo* with Hifiasm[3] using the "-l0" option. Each circular genome was manually retrieved. To reduce calculation times for modification identification, low-complexity regions were masked using Komplexity (https://github.com/eclarke/komplexity) with "--mask --threshold 0.65" options. HiFi reads were mapped to the masked genomes using pbmm2, an official wrapper software for minimap2[4], to calculate interpulse duration ratios. Modification detection and motif prediction were performed using ipdSummary and MotifMaker, respectively, through the SMRT Link package v12.0, with default settings. To account for the detection power of modified nucleotides by SMRT sequencing, m4C and m6A motifs with scores >100000 and all m5C motifs were retrieved as candidate methylated motifs. Motifs with ambiguous sequences were curated manually. For example, TCG**A**VV (where V = A/C/G) was detected in *H. pylori* 3401T, but the spurious partial sequence VV was likely due to incomplete detection of the motif and probably represents the palindrome TCG**A**. Data on methylated motifs in *H. pylori* 26695 were retrieved from a previous study[5].

For phage genomes sequenced in CCS mode (K2 and K23), subreads containing at least three full-pass subreads per polymerase read and with >99% average base-call accuracy were retained as HiFi reads using the standard PacBio SMRT software package, with default settings. For downstream epigenomic analysis, the HiFi reads were converted to add conventional kinetic information. HiFi reads from K2 were then assembled de novo using Canu[6] with the "-pacbio-hifi genomeSize=30k" option, and a single chromosomal genome was manually retrieved. For comparative genomic analysis, the reference KHP30 genome (AB647160.1) was retrieved from the NCBI RefSeq Database, with sub-reads (K2H) and HiFi reads (K2 and K23) mapped to the reference genome using pbmm2 with sub-reads or HiFi read mode, respectively. For epigenomic analysis, both subreads and HiFi reads from each phage were mapped to the assembled KHP30T genome as described above. Nucleotide modifications were predicted using ipdSummary and MotifMaker from SMRT Link packages (v10.2 and v12.0) for the PacBio Sequel and Sequel II systems, respectively, according to the supported systems of each software version, with the default settings. The algorithm used by MotifMaker employs a statistical test for motif identification and is therefore not applicable to small phage genomes due to an insufficient number of motif sequences for de novo identification of modified motifs[7]. Consequently, we focused on host genomes for motif prediction.

CDSs with >33 amino acids in each genome were predicted using Prodigal[8] with default settings or with the "-meta" option for *H. pylori* and phage genomes, respectively. Genes encoding MTases, REases, and DNA sequence recognition proteins (S subunits) were identified using DIAMOND.[9] Sequences were compared against an experimentally confirmed gold-standard dataset from REBASE[10] (downloaded March 1, 2024), with a cutoff e-value of ≤1E-5. REBASE was used to determine the sequence specificity of each MTase and REase. Phage defense systems were predicted using PADLOC.[11] The ANI was calculated using FastANI.[12] Circos plots were generated with the "interacCircos" R package. [13] Homopolymers with more than nine repeat units within or less than 100 bp upstream of RM system genes were retrieved as SSRs associated with the RM system.
